# Tonic REM sleep EEG components predict better mood, cognition and reduce cortical excitability overnight

**DOI:** 10.64898/2026.02.09.704679

**Authors:** Sara Wong, Kiran K G Ravindran, Henry Hebron, Delia Lucarelli, June Lo, John Groeger, William Wisden, Ines R Violante, Derk-Jan Dijk, Valeria Jaramillo

## Abstract

Rapid Eye Movement (REM) sleep makes up approximately 20% of sleep in the adult human and is altered in psychiatric and neurodegenerative conditions. REM sleep comprises two substates, during which eye movements do (phasic REM) and do not (tonic REM) occur. Tonic REM makes up 70-90% of REM sleep but its role in regulating brain function, mood and cognition remain underexplored. We investigated how seven nights of insufficient sleep (6 h time in bed), compared to sufficient sleep, alter periodic and aperiodic components of the phasic and tonic REM sleep electroencephalography (EEG), in 542 sleep recordings of 36 young adults. Associations between phasic and tonic REM sleep EEG and mood, cognitive performance, and overnight changes in cortical excitability as indexed by ‘1/f’ spectral slopes were assessed. Insufficient sleep predominantly affected tonic REM EEG components, specifically the density of theta, the amplitude, density, and frequency of alpha oscillations and the 1/f slope in the 30 to 45 Hz range. These changes associated with mood and cognitive performances, and with overnight reductions in cortical excitability. These results provide evidence for a role of tonic REM sleep in regulating mood and counteracting cognitive deterioration and excitability changes associated with insufficient sleep.

## Introduction

A lack of adequate sleep, which is widespread [1–4], negatively affects emotional control, mood [5–7] and cognitive functioning [6, 8, 9]. In young healthy adults, rapid eye movement (REM) sleep occurs predominantly at the end of the night, so shortening the duration of the sleep period by waking early or delaying bed time disproportionally reduces REM sleep [10–12]. Reduced REM sleep has detrimental effects on mood [13–15] and cognition [16–18]. However, most studies employing experimental sleep manipulations focus on their effects on NREM sleep, in particular slow-wave activity. This leaves the impact of insufficient sleep on REM sleep and the characteristics of field potentials recorded from the scalp, (i.e. EEG) during this sleep stage underexplored.

REM sleep consists of two distinct substates: phasic REM, which displays bursts of rapid eye movements and muscle twitches, and tonic REM, the longer quiescent portions of REM sleep devoid of eye movements [19]. These substates differ in terms of arousal thresholds, information processing, and cortical activity yet are often examined as a single vigilance state (REM). Comparisons of the EEG during phasic and tonic REM sleep demonstrate higher low frequency (∼2-6 Hz) and lower high frequency (∼10-30 Hz) activity in phasic compared to tonic REM [20, 21]. However, the functional roles of phasic and tonic REM sleep are not well-established. Phasic REM is hypothesized to promote processing of emotional experiences, important for emotion regulation and fear extinction as activation of brain networks related to emotional processing [13, 22–24] is observed during this state. Accordingly, increased rapid eye movement density (i.e. phasic REM) is seen in patients with depression and post-traumatic stress disorder (PTSD), though it remains unclear whether this represents a compensatory or maladaptive change [25–27]. The functional role of tonic REM remains even more elusive. As amplitudes of auditory evoked potentials elicited by sounds are higher and arousal thresholds lower during tonic compared to phasic REM [28–30], it has been hypothesized that tonic REM may allow the sleeping individual to monitor the environment for potential danger in-between phasic episodes [31]. However, it seems unlikely that this is its sole function as this substate extends over 70-90 % of human REM sleep (reports differ likely depending on how phasic and tonic REM sleep are scored e.g. [28, 29, 32]) and it therefore potentially holds undiscovered roles in supporting daytime functions. Given that previous studies have returned discrepancies in REM sleep EEG characteristics between rodent and human studies [33] and that rodent studies have relied on muscle or EEG activity instead of rapid eye movements to score phasic and tonic REM [34], the need for human studies to examine phasic and tonic REM as separate neural states is required.

Within phasic and tonic REM sleep, the brain generates oscillations in the theta (∼2-6 Hz) [35, 36] and alpha (∼6-12 Hz) range [37, 38], but the exact contribution of these oscillations during REM sleep to daytime functions remain to be determined. They have been suggested to promote memory consolidation [39–43] and have been linked to dream recall upon awakening [44–46]. However, in these studies, oscillatory activity was commonly estimated as EEG power in a certain frequency range. Yet, EEG power represents a combination of oscillatory (periodic) and non-oscillatory (aperiodic) background activity [47], the latter of which can confound EEG oscillatory estimates. Aperiodic activity can be estimated by fitting an exponential decay function to the power spectrum, with the exponent (or “1/f” spectral slope) characterising how power decreases as frequencies increase. Periodic activity can then be estimated more accurately by detecting spectral peaks exceeding the 1/f aperiodic background activity above a threshold [48]. Using this approach, both theta and alpha oscillations during REM sleep can be detected [29, 45, 46]. Nevertheless, this method has rarely been used to study the function of these oscillatory activities.

One proposed mechanism by which sleep may exert beneficial effects on brain function after sleep is by the reduction of neuronal excitability. Indeed, cortical excitability increases with time awake and decreases during sleep [49–51]. Excitability changes are thought to occur through changes in neuromodulators such as reduced noradrenaline levels [52], and changes in synaptic strength [53]. In rodents, brain oscillations in the theta range (∼4-11 Hz in rodents) support both synaptic potentiation and depression depending on the timing of neuronal activity relative to the theta phase [54]. Theta oscillations also reduce excitatory neuron activity across the nocturnal sleep period and across NREM-REM-NREM triplets [55, 56]. Similar studies in humans are more limited given the challenges of measuring excitability non-invasively. However, recent studies provided evidence that the spectral slope is a good indicator of network excitation-inhibition balance, with steeper, i.e. more negative, slopes indicating lower asynchronous neuronal activity and higher inhibition [57, 58]. The slope in the high frequency range (∼30-50 Hz) correlates with excitatory/inhibitory neuron activity and tracks changes in excitation/inhibition across vigilance states in mice [58]. Similarly, this slope successfully differentiates between arousal levels across vigilance states in humans as it steepens from wakefulness to NREM to REM sleep and is also steeper under propofol-induced anaesthesia compared to wakefulness [58, 59].

Here, we sought to characterise the periodic and aperiodic components of the phasic and tonic REM sleep EEG and their regulation in young healthy adults by comparing a week of insufficient sleep to a week of sufficient sleep. We also investigated relationships between EEG component changes and changes in mood and cognition.

Finally, we explored associations with changes in NREM spectral slopes from the beginning to the end of the sleep period, as an indicator of sleep-dependent excitability changes. Overall, we found that insufficient sleep primarily affected tonic -rather than phasic-, periodic and aperiodic EEG components. Insufficient sleep increased theta density and steepened spectral slopes, components which positively associated with positive mood, cognition, and excitability reductions.

## Results

### Effects of insufficient sleep on total sleep duration and vigilance states

Thirty-six adults completed an extensive residential in-lab crossover protocol, including adaptation and baseline nights (8 h each), followed by seven-night periods of either sufficient (10 h in bed) or insufficient sleep (6 h in bed) in a randomized order separated by a minimum of 10 days (**Suppl Fig S1**). A total of 542 nights were included in the analyses (see ‘Participants and study protocol’ section for detailed protocol).

Compared to sufficient sleep, all seven nights of insufficient sleep decreased Total Sleep Time (TST) (F=471.03, DF=8), the duration of wake after sleep onset (WASO) (F=31.04, DF=8), NREM (N2 + N3) (F=211.81, DF=8), and REM sleep (F=91.33, DF=8), including both phasic (F=59.79, DF=8) and tonic (F=59.35, DF=8) substates (night*condition p<0.001; **Fig 1a**, **Suppl Fig S2a, Table S1)**.

**Figure 1.**
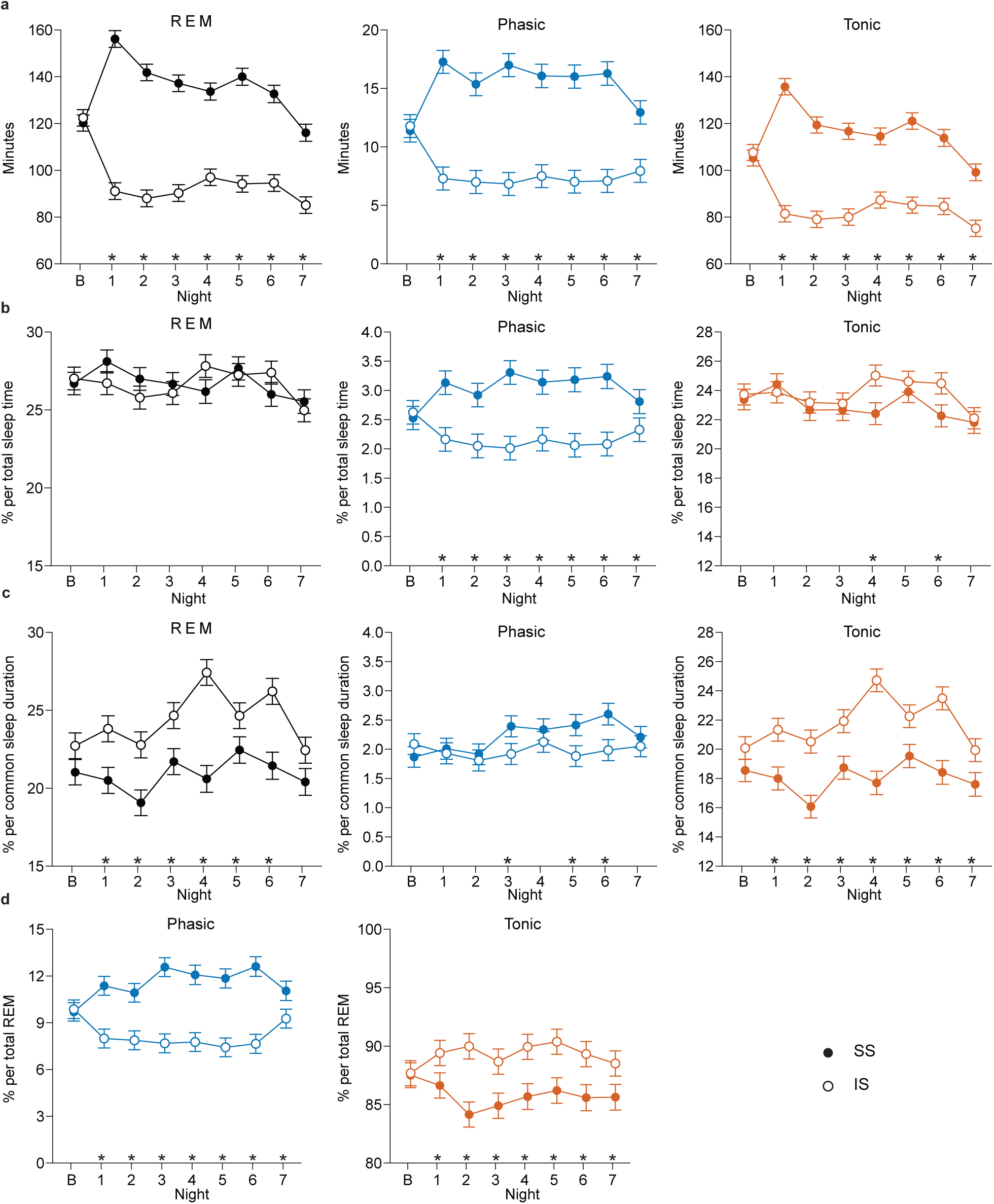
Changes in proportion of REM, phasic and tonic REM sleep during seven nights of insufficient sleep (IS, open circles) compared to sufficient sleep (SS, filled circles) in a. absolute amount (see Supplementary Table S1 for sleep (sub)stage durations for each night) or in b. expressed as percentage of total sleep time, c. as percentage of common sleep duration across SS and IS and across all participants for each respective night, d. as percentage of total REM sleep. Statistical comparisons were made using linear mixed-effects models (lme’s, see Supplementary Table S4 for detailed models) and Tukey post-hoc tests for comparison between conditions for each night (* p < 0.05 – p < 0.001). Data are presented as estimated means resulting from the lme’s averaged across all participants (n = 36) for each night. Error bars indicate the standard error of the mean (s.e.m). REM = rapid eye movement sleep; B = baseline

After accounting for total sleep time, there were no changes in the overall proportion of REM sleep as reported in our previous study [10] (**Fig 1b**). However, when only considering the common sleep duration across both conditions for each respective night (i.e., the data was truncated to include the same sleep duration for SS and IS and across participants which ranged from minimum 262.5 to maximum 337.5 minutes), REM sleep proportion increased across the week of IS (night*condition p<0.001, F=11.73, DF=8; **Fig 1c**) whereas NREM sleep proportion did not show major changes (**Suppl Fig S2c**). This difference suggests an increase in REM sleep at the beginning of the night which gets ‘washed out’ when considering the whole night (higher proportions of REM towards the end of the sleep period will result in higher REM per TST in SS due to the longer sleep period). We next wondered whether there was a change in substate proportions within REM sleep. Indeed, insufficient sleep increased the proportion of tonic REM in the latter days of the week when expressed as % TST and across the week when expressed as % REM (% TST: night* condition p=0.06, F=1.92, DF=8; % REM: night* condition p<0.001, F=10.31, DF=8) and decreased the proportion of phasic REM (% TST: night* condition p<0.001, F=19.88, DF=8; % REM: night* condition p<0.001, F=22.36, DF=8) throughout the week (**Fig 1b and d**). Thus, the overall increase in REM sleep (% common sleep duration) seems to have resulted from a tonic (rather than phasic) REM sleep rebound.

### Phasic and tonic REM sleep EEG periodic and aperiodic components under baseline conditions

When examining the EEG, we confirmed previously reported differences in power spectral density between phasic and tonic REM sleep under baseline conditions: phasic REM exhibited higher spectral power in 1-6 Hz and 32-45 Hz range whereas tonic REM showed higher spectral power in the 12-21 Hz range [20] (**Fig 2a**).

**Figure 2.**
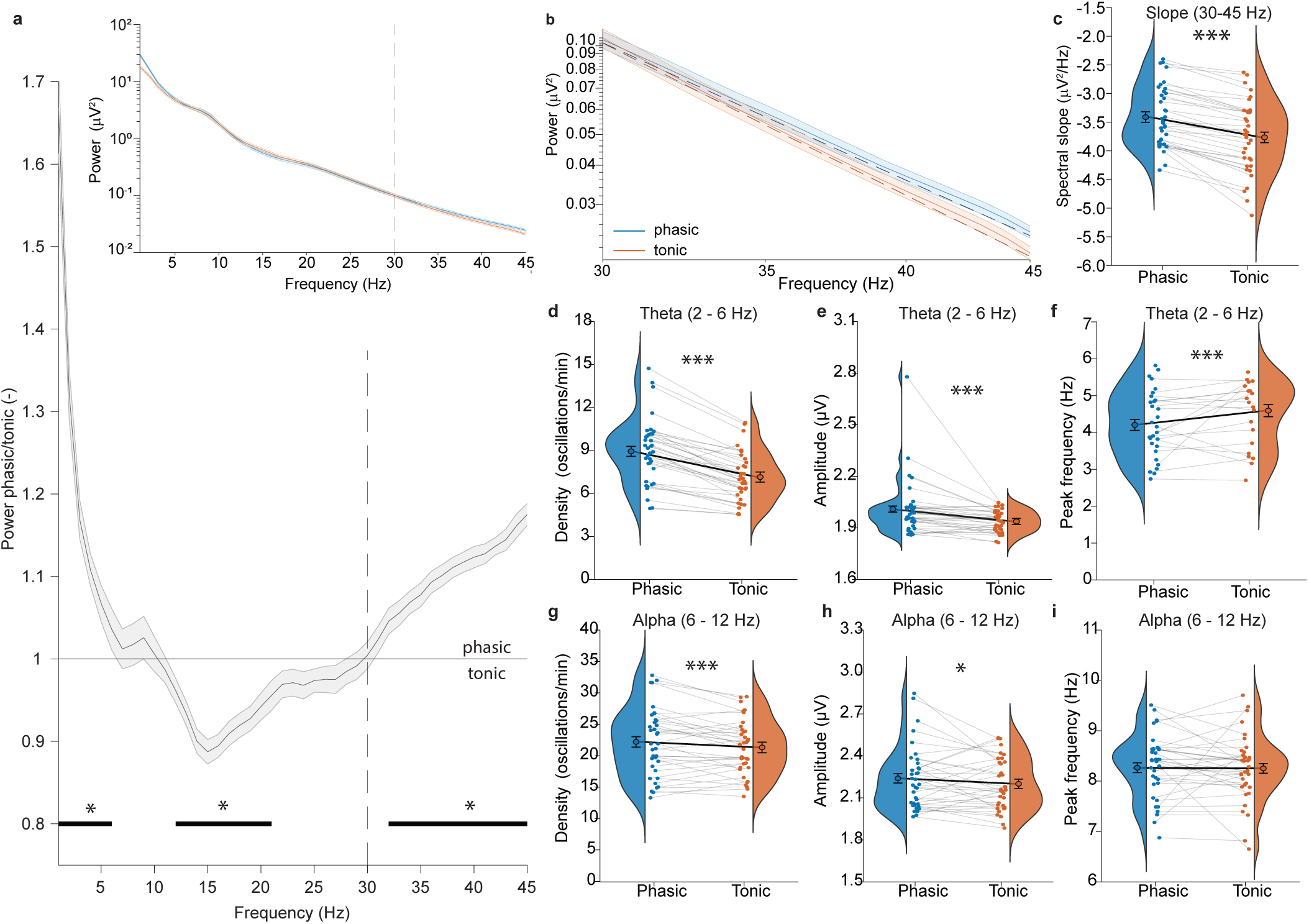
Differences in phasic versus tonic REM sleep power spectra, periodic and aperiodic EEG characteristics during baseline nights. a. Phasic and tonic REM sleep EEG power spectra for 1-45 Hz, plotted as phasic/tonic ratio (major panel) and as absolute spectra on a log-log scale (insert) and b. REM EEG power spectra in the 30 – 45 Hz range, plotted on a log-log scale. Power spectra are represented by solid lines and aperiodic fits by dashed lines. Power spectra and fits were averaged across participants (n = 36) and electrodes. Shaded area represents the standard error of the mean (s.e.m.) across participants for the power spectra. c. Phasic versus tonic REM EEG spectral slope in the 30 – 45 Hz range. d-f. Phasic versus tonic theta (2-6 Hz) oscillations’ density, amplitude and peak frequency. g-i. Phasic versus tonic alpha (6-12 Hz) oscillations’ density, amplitude and peak frequency. Statistical comparisons between phasic and tonic REM sleep were made using linear mixed-effects models and anova (see Supplementary Table S4 for detailed models, * p<0.01 – p<0.001). Coloured circles indicate average values of SS and IS baseline nights for each participant and violins show their distributions. Estimated means resulting from the lme’s averaged across participants (n = 36) are indicated with open black circles. Error bars indicate the s.e.m. Grey and black lines connect the data points from the same participant and average, respectively.

Using the Spectral Parameterization Resolved in Time (SPRiNT) algorithm [60], we performed aperiodic fits and obtained spectral slopes in the high frequency range (30-45 Hz, **Fig 2b**, see **Suppl Fig S3** for results in the 1-30 Hz range) and replicated previously observed differences across vigilance states: spectral slopes were steepest in REM, followed by NREM and wakefulness (**Suppl Fig S4a**). A comparison between phasic and tonic REM sleep revealed that tonic REM was associated with steeper spectral slopes (p<0.001, F=94.93, DF=1; **Fig 2c**) across all scalp regions (**Suppl Fig S4b**). Thus, phasic and tonic REM sleep differ in aperiodic activity, the steeper slopes in tonic REM indicating reduced excitation and/or stronger inhibition in this substate.

We next sought to characterise periodic EEG components of phasic and tonic REM sleep using the extended Better Oscillation detection (eBOSC) method [48]. Oscillations were detected in the 1-30 Hz frequency range and assigned to phasic or tonic REM sleep based on whether they occurred in a phasic/tonic 1-s segment (see methods for details). As 2-6 Hz oscillation density was highest in frontocentral regions in REM sleep (**Suppl Fig S5a**) which fit the description of “sawtooth waves” [36, 61, 62] and 6-12 Hz oscillation density was highest in occipital regions during wakefulness followed by REM, (**Suppl Fig S5c**) which is typical for alpha oscillations [63] we refer to these oscillations as “theta” and “alpha” from here for simplicity. When comparing phasic and tonic REM sleep, phasic REM exhibited higher density and amplitude of theta (density: p<0.001. F=136.16, DF=1; amplitude: p<0.001, F=28.18, DF=1) and alpha oscillations (density: p<0.001, F=13.51, DF=1, amplitude: p<0.05, F=4.94, DF=1) (**Fig2d, e, g and h**), in most scalp regions for theta and in occipital regions for alpha (**Suppl Fig S6a, b, d and e**). Tonic REM displayed faster theta individual peak frequency (p<0.001, F=13.53, DF=1; **Fig 2f**) in frontal and occipital regions (**Suppl Fig S6c**) and phasic REM had higher alpha individual peak frequency in frontal regions (**Suppl Fig S6f**). In summary, we confirmed the occurrence of theta and alpha oscillations in REM sleep, and observed higher densities and amplitudes in phasic compared to tonic REM.

### Effects of insufficient sleep on periodic and aperiodic phasic and tonic REM sleep components

We next evaluated changes in phasic and tonic aperiodic and periodic EEG components during the seven nights of sleep restriction (6 hours in bed, i.e. insufficient sleep) compared to sufficient sleep (10 hours in bed). Spectral slopes (night*state*condition p<0.01, F=2.78, DF=8), theta density (night*state*condition p<0.001, F=4.85, DF=8), alpha amplitude (night*state*cond p<0.001, F=3.86, DF=8), alpha density (night*state*condition p<0.001, F=4.70, DF=8), and alpha peak frequency (night*state*condition p<0.05, F=2.16, DF=8) (**Fig 3**) changed depending on condition and substate across the week. Changes occurred primarily in tonic REM, where spectral slopes steepened with insufficient sleep whereas phasic REM slopes became flatter, i.e. became less negative, towards the end of the week (Tukey-adjusted pairwise comparisons, estimate=0.12, SE =0.04, p<0.05 on night 5, **Fig 3a**), indicating increased inhibition/excitation in tonic REM sleep with insufficient sleep, and the opposite for phasic REM. Insufficient sleep progressively increased tonic theta density (Tukey-adjusted pairwise comparisons, p<0.05 - 0.001 on night 1 and 3-7, **Fig 3b**) and decreased alpha amplitude (Tukey-adjusted pairwise comparisons, p<0.01 - 0.001 on night 3-7, **Fig 3c**) and density (Tukey-adjusted pairwise comparisons, p<0.05 - 0.001 on night 2-7, **Fig 3e**). Tonic alpha frequency decreased with IS (Tukey-adjusted pairwise comparisons, p<0.05 - 0.01 on nights 4-5, **Fig 3d**). No changes were found for other oscillation characteristics (**Suppl Fig S7**).

**Figure 3.**
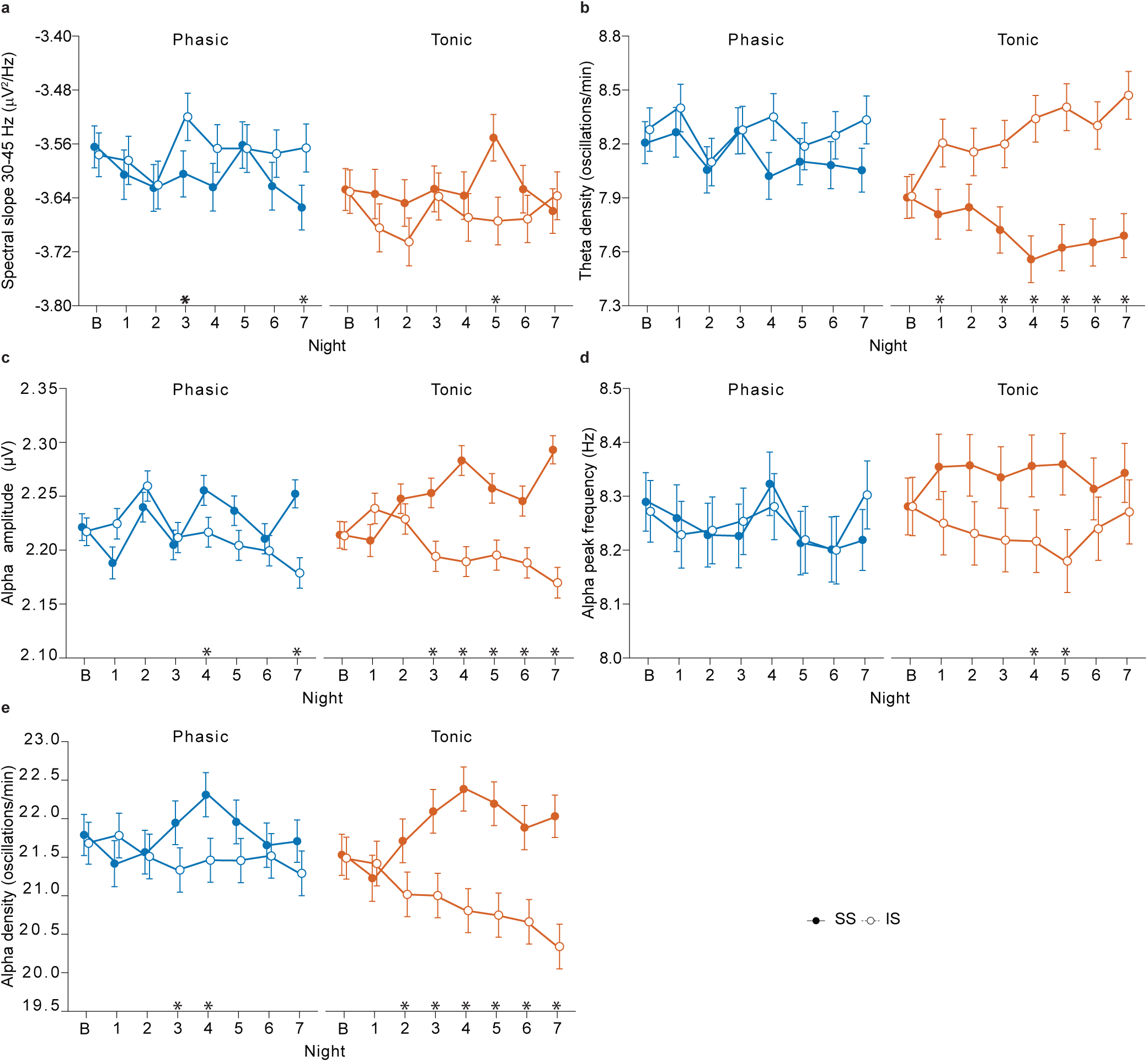
Changes in phasic and tonic oscillatory characteristics and spectral slopes during insufficient sleep (IS, open circles) compared to sufficient sleep (SS, filled circles). a. Changes in phasic and tonic spectral slope in 30 – 45 Hz range. b. Changes in phasic and tonic theta oscillation density. c. Changes in phasic and tonic alpha oscillation amplitude. d. Changes in phasic and tonic alpha oscillation peak frequency. e. Changes in phasic and tonic alpha oscillation density. Statistical comparisons were made using linear mixed-effects models, anova (lme’s, see Supplementary Table S4 for detailed models) and Tukey post-hoc tests for comparison between conditions for each night for the same state (* p<0.05 – p < 0.001). Data are presented as estimated means resulting from the lme’s averaged across participants (n=36) for each night. Error bars indicate the standard error of the mean (s.e.m). B = baseline.

To gain further insight into the regulation of REM sleep substates, periodic and aperiodic characteristics, we also explored within-night dynamics of these changed characteristics (with IS) during baseline nights. The probability of phasic REM sleep increased from the first to the second REM episode and remained stable thereafter (**Suppl Fig S8a**). Spectral slopes tended to become flatter in both REM substates (**Suppl Fig S8b**), and theta density increased towards the end of the night (**Suppl Fig S8c**). Alpha amplitude decreased for both REM substates (**Suppl Fig S8d**). Individual alpha peak frequency increased in tonic REM (**Suppl Fig S8e**) while alpha density increased for phasic REM (**Suppl Fig S8f**).

Overall, tonic REM displayed more pronounced changes in both aperiodic and periodic EEG components in response to insufficient sleep.

### Relationship between changes in phasic and tonic REM EEG components and changes in mood during insufficient sleep

Over the course of the week of the sleep intervention, General Positive Affect (GPA) declined, with a larger reduction for insufficient compared to sufficient sleep (day*condition p<0.001, F=37.08, DF=7, **Fig 4a**). Similarly, General Negative Affect (GNA) increased, however, in this case to a lesser extent with insufficient sleep (day*condition p<0.001, F=13.42, DF=7). These changes indicate a general blunting of emotions with insufficient sleep.

**Figure 4.**
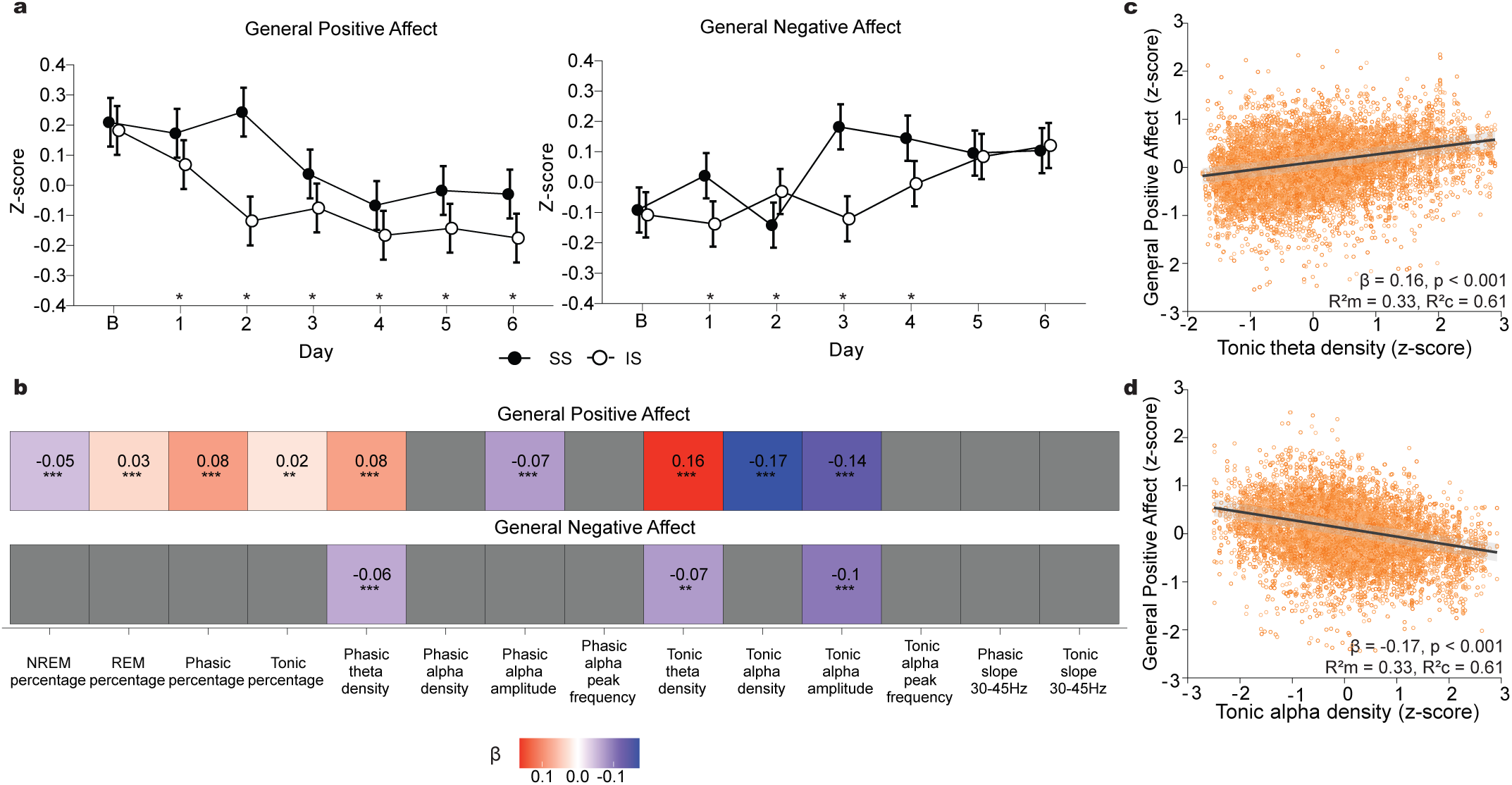
Changes in mood and their relationship with phasic and tonic REM sleep proportion, oscillatory characteristics and spectral slope during sufficient and insufficient sleep. a. Changes in general positive and negative affect during seven nights of insufficient sleep (IS, open circles) compared to sufficient sleep (SS, filled circles). Statistical comparisons were made using linear mixed-effects models (lme’s, see Supplementary Table S4 for detailed models) and Tukey post-hoc tests for comparison between conditions for each night (* p<0.05 – p < 0.01). Data are presented as estimated means resulting from the lme’s averaged across participants (n = 36) for each night. Error bars indicate the standard error of the mean (s.e.m). b. Relationships between general positive and negative affect and vigilance state proportion, oscillatory and aperiodic measures which showed significant differences between SS and IS were assessed using lme’s (data included from both SS and IS, see Supplementary Table S4 for detailed models). Only significant relationships (* p<0.05, ** p<0.01, *** p<0.001, FDR [False-Discovery Rate]-corrected) are coloured and corresponding standardized beta values indicated; red colours indicating positive and blue colours negative relationships. c. Strongest relationships were found with tonic theta density (top) and tonic alpha density (bottom). Black line represents partial linear regression with 95% confidence interval shaded in grey; orange circles represent partial residuals. B = baseline. R2m= R2 marginal, R2c = R2 conditional.

We next evaluated associations between changes in the sleep characteristics that changed with IS from baseline and mood changes from baseline, by controlling for baseline sleep characteristics and mood values [64, 65] (see **Suppl Table S4** and methods for details on statistical model, **Fig 4b**). A stronger association with higher positive affect was found for phasic compared to tonic REM sleep (β=0.08, for phasic REM vs β= 0.02, tonic REM). Conversely, the proportion of NREM sleep was associated with lower positive affect (β= -0.05). However, tonic REM periodic components showed stronger associations with mood: higher tonic theta density correlated most strongly with higher positive affect the following day (β= 0.16, **Fig 4c**) and was also associated with lower negative affect (β= -0.07) indicating a relationship with better mood in general. In contrast, tonic alpha density was related to lower positive affect (β= -0.17) whereas alpha amplitude related with both lower positive and negative affect (β=-0.14 with GPA, β=-0.10 with GNA; **Fig 4b and d**). The associations for theta density and alpha amplitude with GPA were also observed for phasic REM (albeit to a weaker extent than for tonic REM) and were likewise present during the baseline night (**Table S2**). No relationships were found between spectral slope and mood changes.

Taken together, our results show that insufficient sleep decreases both positive and negative affect, in accordance with previous analyses of these data [7]. Moreover, and novel to this study, REM sleep features showed divergent relationships with mood when sleep was insufficient. A decrease in phasic REM sleep was linked to more deterioration of mood, whereas in tonic REM sleep, increases in theta density and decreases in alpha amplitude and density were associated with positive mood changes.

### Changes in daytime cognitive functions with insufficient sleep

To further investigate the contribution of phasic and tonic REM sleep to daytime brain function, we analysed the extensive set of performance measures that were collected five times per day, throughout the protocol. Using a principal component analysis (PCA), we identified four principal components (PCs) that accounted for 46.7 % of the variance across the 46 cognitive measures assessed (**Fig 5a**). PC1 (19.7 % of variance), was named “effort” because the highest loadings were found for ratings of how much effort it took participants to perform the working memory (i.e. N-back) tasks and to stay awake. In addition, performance accuracies across various tasks displayed high negative loadings, indicating that this component also reflects performance inaccuracy. PC2 (12.3 %) was labelled “response time” as response times had the highest loadings onto this component, and high negative loadings were found for performance accuracies in the pursuit tracking task which depends on how quickly participants respond to changes in target trajectory. PC3 was named “response time consistency” (8.3 %) due to the highest negative loadings for the standard deviation of reaction times across various tasks. Finally, PC4 was termed “attentional lapses” (6.4 %) because of the high positive loading for the number of lapses and the high negative loading for the speed of the slowest 10 % of responses in the psychomotor vigilance task, which measures sustained attention.

**Figure 5.**
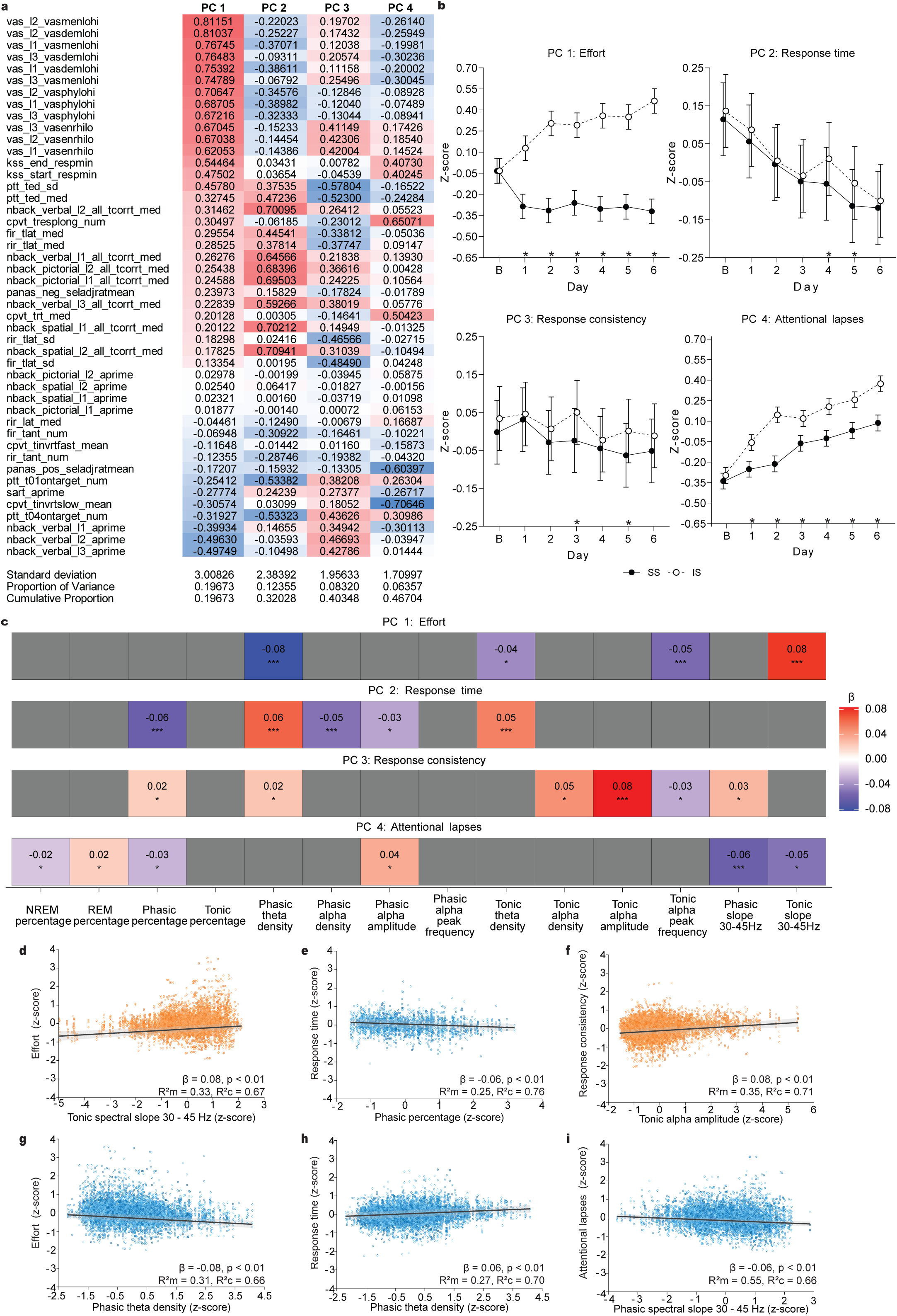
Changes in cognition and their relationship with phasic and tonic REM sleep proportion, oscillatory and aperiodic characteristics during sufficient and insufficient sleep. a. Contribution of cognitive test measures to first four principal components (PCs). Values indicate loading onto the PCs, red colours indicate a positive relationship, blue colours indicate a negative relationship with that component (sorted by loading onto PC1). The standard deviation, proportion of variance and cumulative proportion are shown for each of the four PCs. b. Changes in the four PCs during seven nights of insufficient sleep (IS, open circles) compared to sufficient sleep (filled circles). Statistical comparisons were made using linear mixed-effects models (lme’s, see Supplementary Table S4 for detailed models) and Tukey post-hoc tests between conditions for each night (* p<0.05 – p < 0.001). Data are presented as estimated means resulting from the lme’s averaged across all participants (n=36) for each night. c. Relationships between each PC and vigilance state proportion, oscillatory and aperiodic measures which showed significant differences between SS and IS, were assessed using lme’s (data included from both SS and IS, see Supplementary Table S4 for detailed models). Only significant relationships (* p<0.05, ** p<0.01, *** p<0.001, FDR [False-Discovery Rate]-corrected) are coloured and corresponding standardized beta values indicated; red colours indicating positive and blue colours negative relationships. d-i. Strongest relationships of PC1-4 were found with phasic and tonic spectral slope (30 – 45 Hz), phasic theta density, phasic REM sleep percentage and tonic alpha amplitude. Black line represents partial linear regression with 95% confidence interval shaded in grey, orange and blue circles represent partial residuals. B = baseline. R2m= R2 marginal, R2c = R2 conditional.

Over the week all four PCs changed significantly (for all PCs: day*cond p<0.01, DF=7, PC1: F=459.03, PC2: F=3.11, PC3: F=3.72, PC4: F=64.82, **Fig 5b**). Insufficient sleep increased “effort” progressively throughout the week, showing increased perceived cognitive effort and reduced performance accuracies. “Response time” decreased for both sleep conditions, indicating faster responses over time, likely due to learning effects. For PC2, differences between sleep conditions were only observed on days four and five in which response times were slower with insufficient sleep. Changes in “response time consistency” were subtle towards the end of the week, with more consistent responses for insufficient sleep (day 3 and 5). Lastly, “attentional lapses” increased across the week for both sleep conditions, which was more pronounced with insufficient sleep.

### Relationship between changes in phasic and tonic REM EEG components and changes in daytime cognitive functions during insufficient sleep

Next, we evaluated associations between periodic and aperiodic EEG changes with insufficient sleep and changes in the PCs of cognition.

Focusing on the strongest relationships for each of the PCs, steeper tonic spectral slopes (i.e. more negative) and higher phasic theta density were associated with less perceived effort and better performance (PC1 - tonic slope 30-45 Hz: β=0.08, phasic theta density, β=-0.08, **Fig 5c, d and g**). Higher phasic percentage and lower phasic theta density predicted faster “response times” (PC2 - phasic percentage β= -0.06, phasic theta density β = 0.06, **Fig 5c, e and h**). Higher tonic alpha amplitude correlated with more consistent responses (PC3 β = 0.08). Finally, steeper phasic spectral slopes were linked to an increased number of attentional lapses (PC4 β = -0.06). Most of these relationships were not observed during baseline night except the relationship between phasic theta density and “response time” (**Table S3**).

In summary, steeper tonic spectral slopes and increased phasic theta density were most strongly linked with better cognitive performance in the domain that explained most variance in the cognitive data and showed the strongest deterioration with insufficient sleep (PC1).

### Relationship between phasic and tonic REM EEG components and overnight reduction in excitability levels

REM sleep has been proposed to promote a reduction in excitability levels across the night, which has been suggested to depend on high inhibitory activity [33]. We therefore explored whether spectral slopes during phasic and tonic REM sleep were associated with overnight changes in NREM spectral slopes, as an indicator of excitability reductions. We observed a NREM slope steepening from the first to the last NREM episode of the night indicating an overnight reduction in excitability, as in [33], for both the sufficient and insufficient sleep condition (for both SS and IS: time p<0.001, DF=1, SS: F=233.30, IS: F=56.09, **Fig 6a and b**). We also extracted successive NREM-REM-NREM sleep periods (thereafter referred to as triplets) to examine changes in spectral slopes from the NREM period preceding REM to the NREM period following REM and observed a similar steepening of NREM slopes from pre- to post-REM sleep periods (**Suppl Fig S9a).** In contrast to NREM, phasic and tonic spectral slopes did not change (**Fig 6d, g and h**) or flattened across the night (flattening during phasic REM for IS p<0.001, F= 18.88, DF=1, **Fig 6e**). When comparing between the conditions, overnight NREM slope steepening was reduced with insufficient sleep (night*condition p<0.001 F=6.17, DF=8, **Fig 6c**), which was also the case across NREM-REM-NREM triplets (night*condition p<0.001, F=15.08, DF=8, **Suppl Fig S9g**). While slopes steepened overnight more in tonic REM with insufficient sleep compared to sufficient sleep (night*condition p<0.001, F=3.49, DF=8, **Fig 6i**), no changes were observed for phasic REM with insufficient sleep (**Fig 6f**). Most importantly, both phasic and tonic spectral slopes correlated with a reduction in excitability across the night, yet this association was stronger for tonic REM (phasic: β=0.17; tonic β=0.29, **Fig 6j and k**). This was also the case across NREM-REM-NREM triplets (**Suppl Fig S9b and h**). Moreover, we confirmed overnight changes in spectral slopes, as well as the relationships between phasic and tonic spectral slopes and overnight NREM slope steepening in baseline nights (**Suppl Fig 10**).

**Figure 6.**
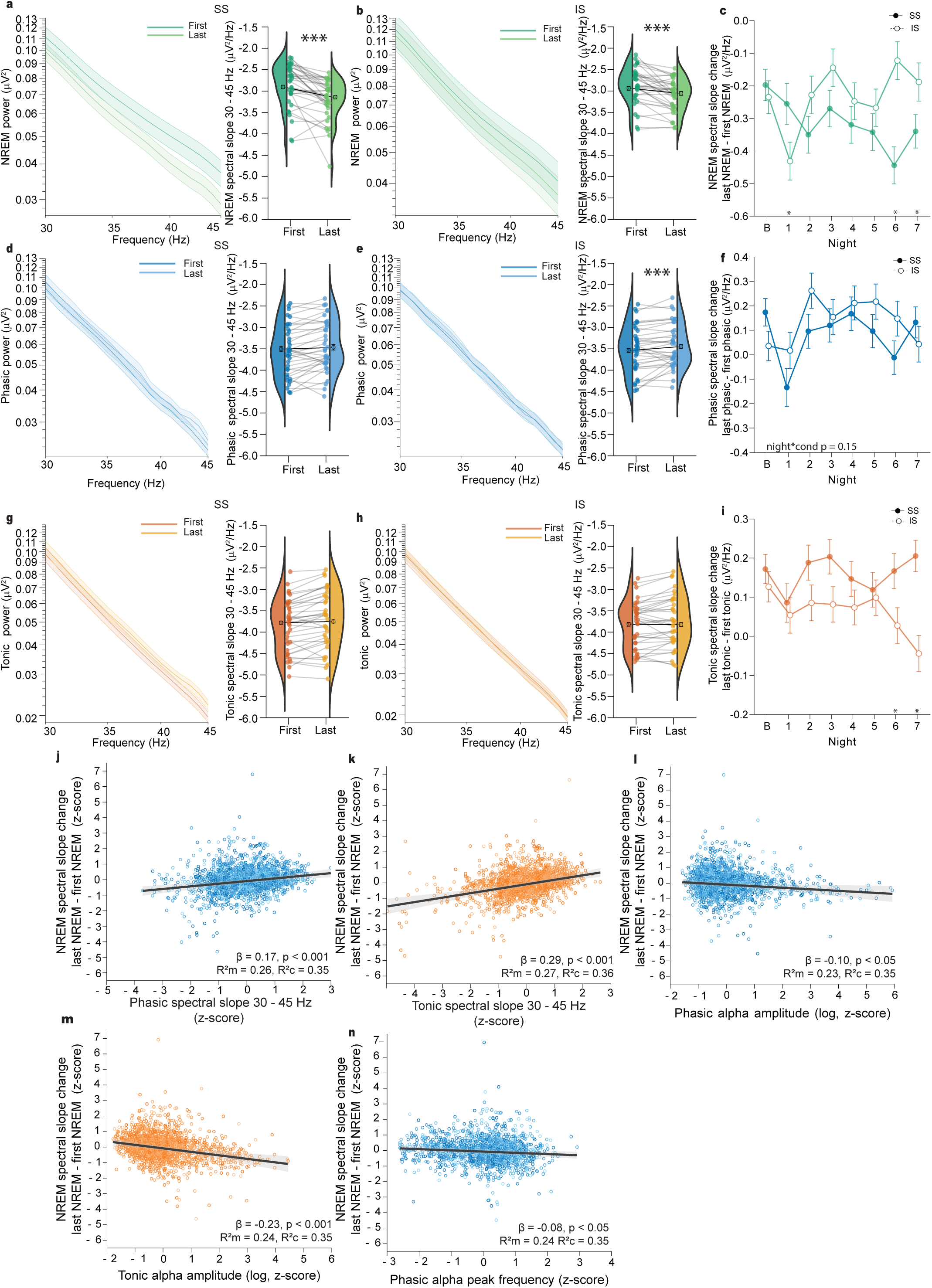
Changes in NREM spectral slopes overnight and their relationship with phasic and tonic spectral slopes, alpha amplitude and theta density. a-b. Changes in NREM spectral slopes in the 30-45 Hz range from the first to the last sleep cycle, during sufficient sleep (SS in a) and insufficient sleep (IS in b). c. Comparison of NREM spectral slope changes from the first to the last sleep cycle (last-first) between SS and IS. d-f. Same as a-c, but for phasic REM. g-i . Same as a-c, but for tonic REM. Statistical comparisons were made using linear mixed-effects models, anova (lme’s, see Supplementary Table S4 for detailed models) and Tukey’s post-hoc comparison between conditions for each night (* p<0.05 - p<0.001). Coloured circles indicate average values of SS and IS nights for each participant and violins indicate their distributions. Estimated means resulting from the lme’s averaged across participants (n = 36) are indicated with open black circles. Error bars indicate the s.e.m. Grey and black lines connect the data points from the same participant and average, respectively. j. Partial linear regression plot showing relationship between NREM spectral slope change overnight with mean phasic spectral slope. k. Partial linear regression plot showing relationship between NREM spectral slope change overnight with mean tonic spectral slope. l. Partial linear regression plot showing relationship between NREM spectral slope change overnight with phasic alpha amplitude. m. Partial linear regression plot showing relationship between NREM spectral slope change overnight with tonic alpha amplitude. n. Partial linear regression plot showing relationship between NREM spectral slope change overnight with phasic alpha peak frequency. Black line represents partial linear regression with 95% confidence interval shaded in grey, orange or blue circles represent partial residuals. All correlations were FDR [False-Discovery Rate]-corrected.

Given that in rodents REM sleep theta oscillations have been linked with synaptic plasticity processes and overnight excitability reductions [54–56], we explored relationships between overnight NREM spectral slope changes and EEG oscillations in REM sleep. Specifically, these were evaluated for theta density and alpha density, amplitude and peak frequency, since these characteristics showed changes with insufficient sleep and it is unclear which frequency range (alpha or theta) represents the human functional analogue to the rodent theta oscillation [66]. Interestingly, we found that larger alpha amplitudes during both phasic and tonic REM sleep associated with larger excitability reductions, again with a stronger relationship for tonic REM sleep (phasic: β = -0.10; tonic: β = -0.23, **Fig 6l and m)**.This relationship was also seen across NREM-REM-NREM triplets for tonic but not phasic REM (**Suppl Fig S9d and j**) and for both REM substates during baseline nights (**Suppl Fig S11d and i**). In addition to alpha amplitude, higher alpha peak frequency during phasic REM was also linked to larger overnight excitability reductions (β = -0.08, **Fig 6n**) whereas no relationship was found with alpha density (**Suppl Fig S12c and d**). Conversely, higher phasic and tonic theta density associated with decreased NREM spectral slope steepening across triplets (SS and IS: **Suppl Fig S9c and I,** baseline**: Suppl Fig S11c and h**), however, no association with theta density was present for overnight NREM slope changes (**Supp Fig12 a and b**).

These results indicate that steep spectral slopes (i.e. low REM sleep aperiodic activity reflecting high inhibitory activity), as well as alpha oscillations may contribute to overnight excitability reductions.

## Discussion

By differentiating between phasic and tonic REM substates and separating oscillatory from aperiodic activity in each substate’s EEG, we found that insufficient sleep resulted in a tonic REM sleep rebound accompanied by pronounced EEG changes. These changes correlated with mood and cognition, suggesting that tonic REM may serve previously unrecognized, fundamental roles in daytime functions.

Among the various aspects of daytime function, mood seems to be promoted by REM theta oscillations, considering we observed the strongest associations between phasic and tonic theta density and positive affect. Although, it should be noted that positive affect reflects higher vigour/energy levels rather than “happiness” per se [7]. Higher theta density during phasic REM and stronger relationship of phasic REM proportion with positive affect are in line with an important role for phasic REM in mood regulation. However, tonic REM may play a role in counteracting mood changes when phasic REM is shortened by insufficient sleep, through increasing its theta density, alongside a stronger relationship of tonic theta density with mood. This result aligns with previous sleep restriction studies reporting an increase in theta power during REM sleep, although they did not differentiate between phasic and tonic REM nor between periodic and aperiodic EEG components [67]. Our observations are also consistent with studies indicating a role for REM sleep theta oscillations in emotional regulation although most studies were conducted in rodents [34, 68] and it remains unclear which frequency range represents the functional analogue in humans. We here provide novel evidence for a potentially similar role of human oscillations in the 2-6 Hz frequency range in emotional regulation. Interestingly, sleep restriction is well-known to have positive effects on depressive symptoms in insomnia [69], and ketamine, which is used in treatment-resistant depression (TRD), reduces phasic REM during the first REM sleep period in TRD [70, 71]. Depression is associated with increased high frequency (alpha and higher) and reduced low frequency (theta) activity during REM sleep [72]. Considering our results, it is plausible that the positive effects of sleep restriction on mood may result from an increase in tonic theta density (and reduction in alpha density/amplitude, see below) following short sleep. It remains to be confirmed whether similar tonic REM sleep EEG changes are present as a consequence of sleep restriction in these clinical populations.

An important new finding is the relationship between steep tonic spectral slopes and reduced “effort”. During insufficient sleep, the spectral slope steepened in tonic REM, indicating that spectral slopes are regulated depending on REM sleep history and may help to counteract cognitive deterioration. A potential mechanism by which steep spectral slopes (indicating increased inhibition [57, 58]) could promote cognition is via a reduction in excitability across the night. Accordingly, we found associations between phasic and tonic spectral slopes and an indicator of overnight excitability changes (i.e. NREM spectral slope changes overnight). In addition to replicate a previously reported relationship between REM sleep and overnight downregulation of aperiodic activity [33], we further elucidate the asymmetrical contribution of REM substates. Our results indicate that tonic REM may be the more important REM substate in reducing excitability and cognitive deterioration with insufficient sleep, considering the steeper slopes under baseline conditions, the spectral slope changes with insufficient sleep, and the relationship of tonic but not phasic, spectral slopes with “effort”. This is relevant in the context of ageing and dementia, which is associated with increased cortical excitability thought to underlie neurodegeneration and cognitive deterioration [73–76].

In addition, given the relationship between theta density and reduced “effort”, the increase in theta density during tonic REM sleep with insufficient sleep may not only help to alleviate mood but also cognitive changes with insufficient sleep. Yet, theta density was not linked to overnight excitability reductions and thus may support cognition via a distinct mechanism. In contrast to REM sleep theta oscillations, alpha oscillations have received less attention, although they occur abundantly in REM sleep in healthy young individuals, as we showed here and as previously reported [37, 38]. Yet, the function of these oscillations remains mysterious. The correlation of alpha density/amplitude with more deterioration of mood here indicates a negative effect of these oscillations on mood. In line with this, increased alpha activity during REM sleep has previously been linked to heightened arousal and sleep state misperception – the subjective impression of being awake while the EEG looks like sleep, which often occurs in individuals with insomnia who frequently suffer from mood disturbances [77–79]. Of note, the changes in alpha density and amplitude in response to insufficient sleep being opposite to those of theta density is reminiscent of the behaviour of these two frequency ranges during wakefulness. During wakefulness, alpha activity is well-known to inhibit theta activity [80–82]. Thus, it is conceivable that the increase in tonic theta density in our study may have resulted from a disinhibition by a reduction in tonic alpha density/amplitude, and that the relationship of alpha with less positive mood may result indirectly i.e. from the inhibitory effect on theta – yet this remains speculative and requires further investigation. Although we only observed a small albeit significant relationship of higher tonic alpha peak frequency with less “effort” it is possible that alpha oscillations may instead contribute to cognitive functions. Interestingly, our finding that phasic and tonic alpha amplitude and phasic alpha peak frequency were also associated with an overnight reduction in our excitability indicator, suggests a potential role of these oscillations in modulating overall excitatory-inhibitory balance. In support of these speculations, REM sleep EEG slowing, indicative of alpha frequency slowing, is associated with cognitive impairment in ageing and dementia [83] [84].

Our study had several limitations. To analyse all 542 nights, we used an automatic classifier based on our own set criteria as there is a lack of consensus in the field on how to define and score phasic and tonic REM sleep [85]. In doing so, we cannot exclude that using slightly different criteria could have affected the results. Furthermore, considering the higher proportion of tonic compared to phasic REM, the estimates for specific EEG features may have been less robust for phasic compared to tonic REM. Nevertheless, our approach had strong agreement with the existing literature in terms of phasic vs tonic spectral power differences and allowed for high-temporal resolution REM substate classification with reduced potential bias associated with manual scoring [29, 86].

In conclusion, our study is the first and largest dataset to separate REM sleep into its phasic and tonic substates as well as its EEG into periodic and aperiodic components to investigate the functional relevance of these substates and components in terms of mood and cognition. Our findings suggest that increases in theta oscillation density and steepening of spectral slopes (i.e. high inhibitory activity) during tonic REM counteract mood and cognitive deterioration with insufficient sleep. Overall, our study provides compelling evidence suggesting that tonic REM is not merely important to provide windows of increased arousal but also to promote mood and cognitive function.

## Materials and methods

### Participants and study protocol

Thirty-six young, healthy individuals (18 males and 18 females; Mean age = 27.6, SD = 4.0 years) completed a single-centre, crossover design sleep manipulation protocol during which they were resident at the Surrey Sleep Research Centre. The research protocol was approved by the University of Surrey’s Ethics Committee, and all participants provided their written informed consent to participate in this study. The data from this study has been published previously [10]. Participants underwent an adaption night (8 h), a baseline night (B, 8 h), seven nights of an extended sleep opportunity (SS, 10 h in bed) which allows for sufficient sleep to maintain performance [8, 87] or sleep restriction (IS, 6 h in bed) i.e. insufficient sleep, followed by total sleep deprivation (SD) of 41 h or 39 h under constant routine and a recovery period of 12 h sleep opportunity. To shorten or extend sleep opportunity, each participant’s habitual sleep midpoint was used to define an individualized 8-hour sleep window, extending 4 hours before and after the midpoint. Insufficient sleep was implemented by reducing 1 hour from both the beginning and end of this window, whereas sufficient sleep was achieved by adding 1 hour to both ends. The order of the two conditions was randomized across participants, and the two sessions were separated by a minimum of 10 days. During the awake periods, participants completed a battery of cognitive tasks, including the Karolinska Sleepiness Scale (KSS), Karolinska Drowsiness Test (KDT), the Psychomotor Vigilance Task (PVT), Sustained Attention Response Task (SART), spatial 1- and 2-back, pictorial 1- and 2-back, integrated 1- and 2-back, Verbal 1-, 2-, and 3-back, Visual Analogue Scale (VAS), Pursuit Tracking Task (PTT), Fixed Interval Repetition (FIR) and Random Interval Repetition (RIR), and Positive and Negative Affect Schedule (PANAS) at five equally spaced opportunities during the SS/IS week and every two hours during SD (**Suppl Fig S1**). Due to differences between the constant routine protocol and the protocol during the first 6 days of IS/SS, the constant routine assessments and recovery night after SD were not included in the analyses. A more detailed description of the participant and methodological criteria is summarised in the original report [8] with detailed changes in accompanying cognitive tasks and PANAS reported by [7, 8] and SWA in [10].

A total of 542 nights were included in subsequent analyses with 34 nights excluded due to participant withdrawal (11 nights), missing data (2 nights) or files with missing/inaccurate lights off/on markers (21 nights).

### Polysomnography acquisition and EEG preprocessing

Polysomnography measures were recorded using Siesta 802 devices (Compumedics Ltd., Abbotsford, Victoria, Australia). Measures included electroencephalogram (EEG; F3, F4, C3, C4, P3, P4, O1, O2 referenced to contralateral mastoids), electromyogram (EMG), electrooculogram (EOG), and electrocardiogram (ECG). EEG electrodes were placed using a standardized 10-20 placement system. A sampling rate of 256 Hz was applied.

EEG data were high-pass filtered at 0.3 Hz, low-pass filtered at 70 Hz and notch filtered at 50 Hz. Sleep data were visually scored by sleep experts in 30-s epochs using Rechtschaffen and Kales criteria [88]. NREM sleep epochs were later updated to fit standard AASM criteria, where sleep stage 1 was converted to N1, sleep stage 2 to N2 and sleep stage 3 plus sleep stage 4 to N3.

### Phasic and tonic REM detection and power analyses

A custom-made automatic algorithm was applied to identify phasic REM, tonic REM, and artefacts per second within each 30-s REM sleep epoch. To achieve accurate differentiation of phasic, tonic, and artefact segments, we implemented a soft voting approach that uses five distinct classifiers (different random forest models and boosting ensembles). The individual classifiers used for the soft voting were trained and validated using 52 sleep recordings from 16 participants. The held-out test dataset comprised of 32 recordings. The labelling of the 1 s segments with each REM epoch in the recordings were performed using an initial version of the manual annotation tool, ScoreREM [89]. Of the total 1 s segments within each REM epoch scored in the 52 recordings, the distribution of the labels was 90.5% Tonic, 7.9% Phasic and 1.5% Artefact. To address this class imbalance during training, we used Synthetic minority over-sampling technique (SMOTE). The models used 28 features (including statical, spectral and nonlinear measures) that were extracted from both EOG electrodes and concatenated. The final classification decision was achieved by summing the probabilities predicted by each individual model across all classes and selecting the class with the highest cumulative probability. This approach balanced out the model biases thus mitigating the overfitting problem formerly seen in the individual models and achieved robust performance on the holdout test data. Our approach provided substantial agreement on the held-out test set with average overall Kappa of 0.66±0.08 (Phasic: 0.67±0.05; Tonic: 0.67±0.08; Artefact: 0.65±0.15).

All analyses included in this manuscript were performed using the labels from the custom-made automatic algorithm. All EEG analyses were performed in Matlab 2021a.

An automatic Independent Component Analysis (AMICA, EEGLAB) was applied to extracted REM segments to remove ocular, movement, or cardiac artefacts. Power was calculated in 1 s epochs (Hann window, 0-45 Hz, in 1 Hz steps, 50 % overlap, averaging 5 windows) using SPRiNT_stft from the SPRiNT toolbox [60]. Absolute spectral power values in the frontal (F3 + F4), central (C3, C4), posterior (P3 + P4) and occipital (O1, O2) regions were averaged.

### Aperiodic background activity estimation

Aperiodic background activity was estimated using Spectral Parameterization Resolved in Time (SPRiNT) [60]. Compared to other aperiodic estimation methods (including eBOSC), SPRiNT was specifically developed to provide estimates of aperiodic activity in a time-resolved manner allowing for estimation of aperiodic activity using very short epoch durations (∼1 s). Briefly, aperiodic parameters (exponent and offset) were estimated using *specparam* resulting from moving short-time Fourier transforms derived power spectral density (see above). This procedure was performed for two frequency ranges: 1-30 Hz and 30-45 Hz. While the lower frequency range represents the range used for oscillation detection, the higher frequency range has been suggested to indicate excitation/inhibition balance and to discriminate better between arousal levels [58, 90]. Furthermore, the 30 – 45 Hz range is less influenced by low-frequency brain oscillations, making it easier to fit aperiodic background activity [57, 91]. Since power spectra were assumed to follow a 1/f power law, 1 s epochs with exponents below 0.2 were excluded from the analysis.

### Oscillation detection

Oscillations across wake, NREM, REM, phasic and tonic REM sleep were detected using the extended Better Oscillation detection (eBOSC) algorithm which has been optimized for time-varying, intermediate-length (∼20-40 s) epoch durations [48], enabling detection of slower oscillations such as delta and theta more easily compared to algorithms using shorter epoch durations (including SPRiNT). eBOSC detects oscillations based on wavelet-derived power exceeding set power thresholds of a specific minimum duration. Oscillations were detected in 30 s epochs (1-30 Hz, 0.25 Hz steps, 95% power threshold). Only oscillations with at least 3 cycles and a duration of 300 ms or more were included in the analyses. Visual inspection of the oscillations’ frequency distribution histograms during REM sleep indicated a bimodal distribution within the 2-12 Hz range, with peaks at ∼4 Hz and ∼8 Hz (**Suppl Fig S11**). As the threshold between these distributions was not easy to determine and varied across participants, we subdivided the oscillations detected into broader frequency ranges: 2-6 Hz and 6-12 Hz. To better understand the oscillations detected within the chosen frequency ranges, we examined the spatial distribution and occurrence of these oscillations across vigilance states. 2-6 Hz oscillation density was highest in frontocentral regions in REM sleep and wakefulness, with highest occurrences during REM sleep (state*region p<0.001, **Suppl Fig S5a**). This topographical distribution fits the description of “sawtooth waves” [36, 61] and frontomedial theta during wakefulness [92] and for simplicity, we will refer to these oscillations as “theta”. REM sleep exhibited highest theta peak frequency compared to wakefulness and NREM across all regions except in the occipital region where there were no differences between NREM and REM (p<0.001, **Suppl Fig S5b**). 6-12 Hz density was highest in occipital regions, with greatest occurrences during wakefulness, closely followed by REM sleep (state*region, p<0.001, **Suppl Fig S5c**). This topographical distribution is typical for alpha oscillations, and for simplicity, we will thus call these oscillations “alpha” [63]. Alpha frequencies were highest during wakefulness, increasing towards the back of the scalp (state*region, p<0.001, **Suppl Fig S5d**). REM alpha frequencies were around 1-1.5 Hz slower.

For each participant and frequency range, the peak frequency, density, and amplitude was determined. Density refers to the number of oscillations that occurred per minute while amplitude refers to the power of each oscillation divided by the power of the 1/f background estimate (SNRMean in eBOSC). For peak frequency estimation, the kernel density estimation was used to smooth the frequency distribution before peak detection using the *findpeaks* function. For phasic and tonic REM, oscillations detected during REM sleep were assigned to phasic REM sleep if they contained samples (from oscillation start to end) that occurred within a phasic 1-s segment, if not they were assigned to tonic REM. To avoid a bias towards phasic REM sleep for density estimation by subdividing waves in this manner, density for each 1-s segment was calculated based on the number of oscillations that contained samples within that segment. If an oscillation spanned multiple segments, it was added as 1/number of segments it spanned to the oscillation count. Phasic/tonic densities were then calculated by averaging this number across all phasic/tonic 1-s segments and multiplying by 60.

### Sleep cycle detection

Sleep cycles were detected based on the criteria defined by [93]. In short, NREM periods of at least 15 minutes followed by REM periods of at least 5 minutes were defined as sleep cycles. For the first sleep cycle, no minimum duration was required for the REM episode.

### Statistical methods

All statistical analyses were conducted in R (v4.4.1) using linear mixed-effects (lme) models using *lme4* v1.1.35.5 with participant as a random factor and compound symmetry coefficient structure, and all post-hoc tests using *anova* and *emmeans* with degrees of freedom computed using the asymptotic (z-test) approach. R2 marginal to account for variance explained by fixed effects and R2 conditional to account for variance explained by fixed and random effects were calculated using the *MuMIn* package v1.48.11 [94]. Estimated means were calculated and plotted using the *modelbased* v0.8.8 package.

To reduce dimensionality of the cognitive data a principal component analysis (PCA) was performed across the 46 measured variables, including all five assessments during days following baseline, SE and SR nights. Time points for which data across all variables were missing (1.9 %) were not included in the PCA and were considered as missing in later analyses. Missing data for time points for which data from at least one variable was present (0.3 %) was imputed with the median of the missing variable and of the respective participant. The PCA was then computed using *prcomp* from the *stats* package. The number of principal components to be included was determined by making a Scree plot (plotting the variance explained by the components in descending order) and finding the “elbow” in the curve.

A table of all forementioned models is included in Supplementary Materials (Table S4). For models investigating changes and correlations during baseline, values for both IS and SS conditions were included. For models investigating periodic, spectral slope, mood and cognition changes over the SS/IS week, baseline values were included as a covariate to evaluate changes from baseline [64, 65] and *night:condition* interaction rather than *condition* alone was included as a covariate account for the fact that “baseline” represents a single unit across both SS and IS baseline nights. Similarly, in models investigating correlations during SS/IS week, baseline values and all nights were included as covariates. For correlations, all numeric values were z-scored normalised and models conducted separately for phasic and tonic REM.

## Data availability

All processed data supporting the findings of this study are available on Zenodo: (made available upon publication).

## Code availability

Code for data and statistical analysis are available on GitHub at https://github.com/saraxwong/phasictonicREM. The MATLAB application used for the manual annotation of phasic/tonic sleep [89] is available at https://github.com/KiranKGR/ScoreREMGUI.

## Acknowledgments

This research is supported by the UK Dementia Research Institute by a Cross-Centre UK Dementia Research Institute post-doctoral award (S.W.) through UK DRI Ltd, principally funded by the Medical Research Council (W.W., UKDRI-5004), the UK Dementia Research Institute (V.J., D.J.D. and K.K.G.R., UKDRI-7206), Care Research and Technology Centre at Imperial College, London, and the University of Surrey, Guildford; the Medical Research Council (S.W. and W.W., APP9329); Swiss National Science Foundation (V.J., P2EZP3_199918, P500PB_217827); Wellcome (V.J. 301007/Z/23/Z), the University of Surrey’s Doctoral College Scholarship Award (H.H.); the Biotechnology and Biological Sciences Research Council (BB/S008314/1, I.R.V.). The data acquisition was funded by Grant FA9550-08-1-0080. For the purpose of open access, the author has applied a CC BY public copyright license to any Author Accepted Manuscript version arising from this submission.

## Author contributions

Conceptualization: S.W., V.J., I.R.V., D.J.D.; Data collection and curation: J.CY.L., J.A.G.; Data analysis: S.W., V.J., D.L., H,H., I.R.V., J.CY.L., J.A.G.; Funding acquisition: D.J.D., W.W., J.A.G., V.J., I.R.V.; Software: S.W., V.J., K.K.G.R., H.H.; Supervision: V.J., D.J.D., J.A.G.; Writing—original draft: S.W., V.J.; all authors contributed to the editing of the final draft.

