## Supplementary materials for "Tonic REM sleep EEG components predict better mood, cognition and reduce cortical excitability overnight"

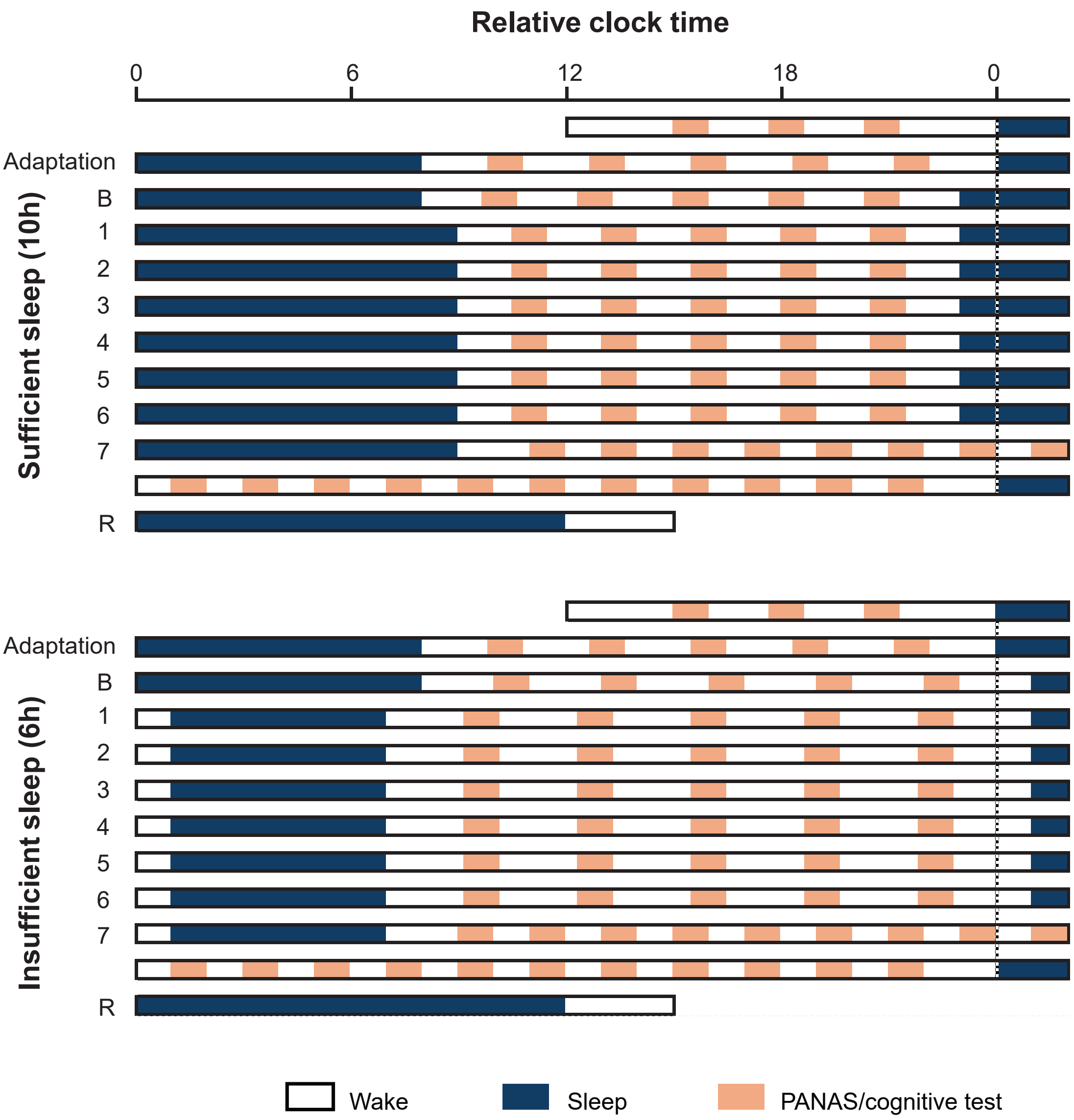


**Supplementary Figure 1**

Study protocol. Each participant underwent a week of sufficient and insufficient sleep in a laboratory setting, in a crossover study with the order of the sleep condition counter-balanced. Each session consisted of an adaptation night (8 hr sleep opportunity) followed by a baseline night (B, 8 hr sleep opportunity) and 7 condition nights (1 – 7, sufficient sleep with 10 hr sleep opportunity; insufficient sleep with 6 hr sleep opportunity), followed by ~ 40 hr of total sleep deprivation under constant routine conditions, and a recovery night (R; 12 hr sleep opportunity). During the awake periods, participants completed a battery of cognitive tasks and Positive and Negative Affect Schedule (PANAS) at five equally spaced opportunities during the SS/IS week and every two hours during SD.

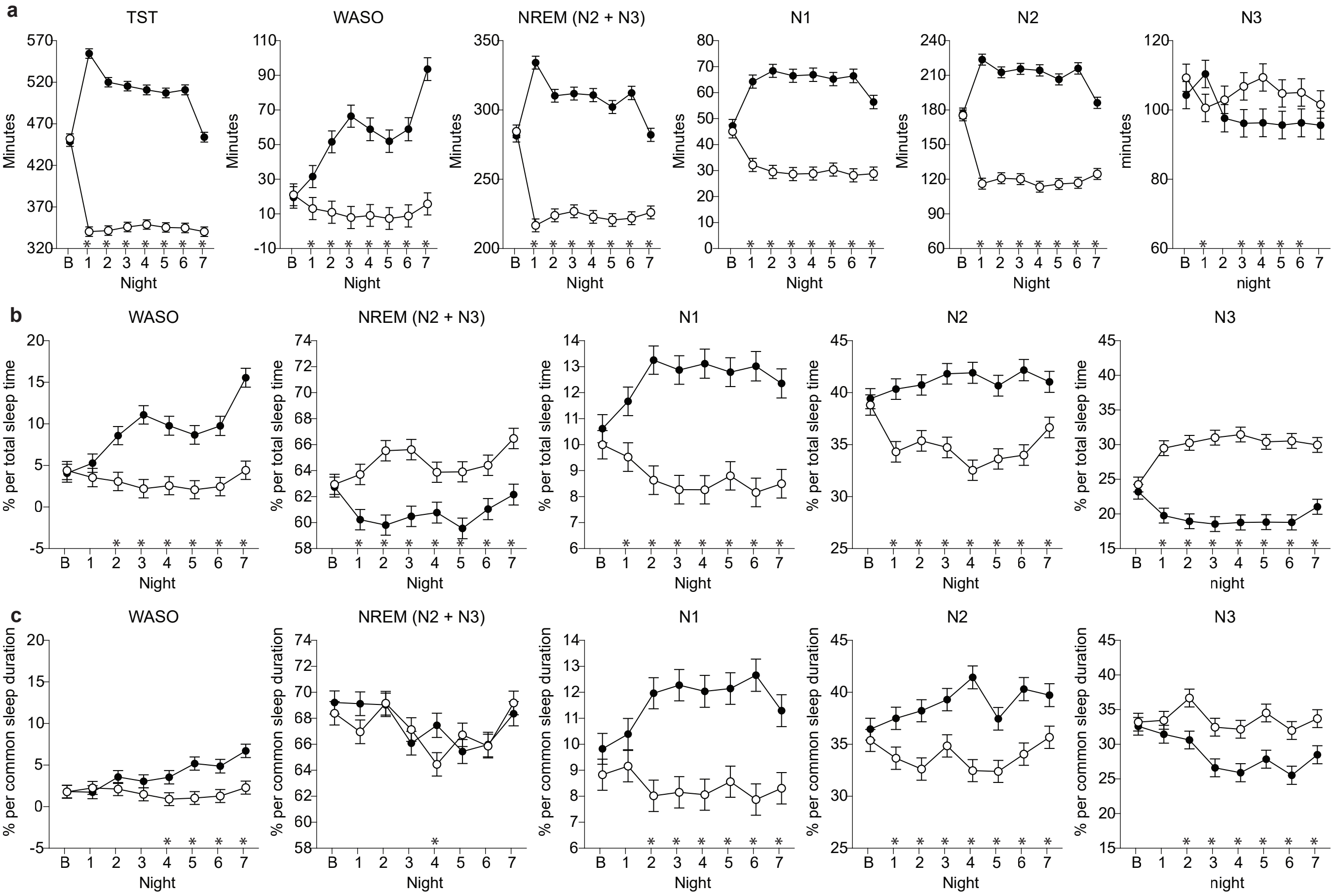


**Supplementary Figure 2**

a. Changes in absolute duration of TST, WASO, NREM, and NREM substages (N1-N3) during seven nights of insufficient sleep (IS, open circles) compared to sufficient sleep (SS, filled circles). b-c. Changes in proportion of WASO, NREM, and NREM substages (N1 – N3) expressed as percentage of (b) total sleep time and (c) common sleep duration across SS and IS and across all participants for each respective night. Statistical comparisons were made using linear mixed-effects models (lme's, see Supplementary Table S4 for detailed models) and Tukey post-hoc tests between conditions for each night (* $p < 0.05$ – $p < 0.001$). Data are presented as estimated means resulting from the lme's averaged across all participants (n = 36) for each night. Error bars indicate the standard error of the mean (s.e.m). TST = total sleep time; WASO = wake after sleep onset; NREM = non-rapid eye movement sleep; REM = rapid eye movement sleep; B = baseline

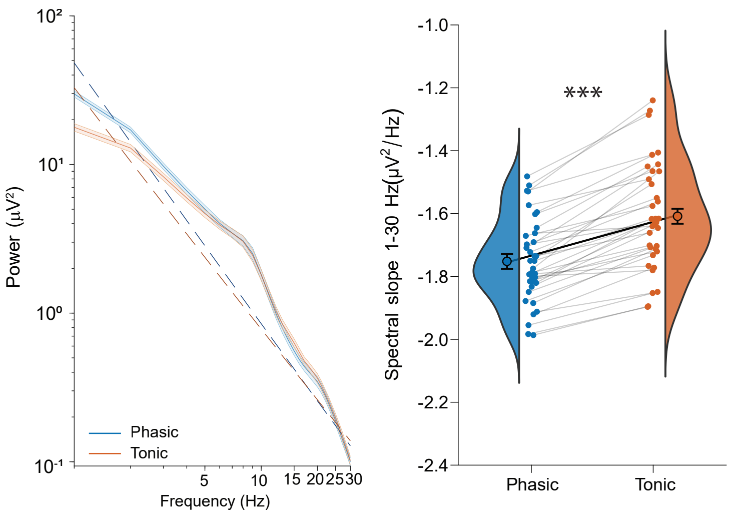


**Supplementary Figure 3**

Differences in phasic versus tonic REM sleep spectral slope in the 1 – 30 Hz range (plotted on a log-log scale) during baseline nights. Power spectra are represented by solid lines, aperiodic fits by dashed lines. Power spectra and fits were averaged across participants (n = 36) and electrodes. Shaded area represents the standard error of the mean (s.e.m.) across participants for the power spectra. Statistical comparisons between phasic and tonic REM sleep were made using linear mixed-effects models (lme's, * $p<0.05$ substage) (see Supplementary Table S4 for detailed models). Coloured circles indicate average values of SS and IS for baseline nights for each participant and violins show their distributions. Estimated means resulting from the lme's averaged across participants (n = 36) are indicated with open black circles. Error bars indicate the s.e.m. Grey and black lines connect the data points from the same participant and average, respectively.

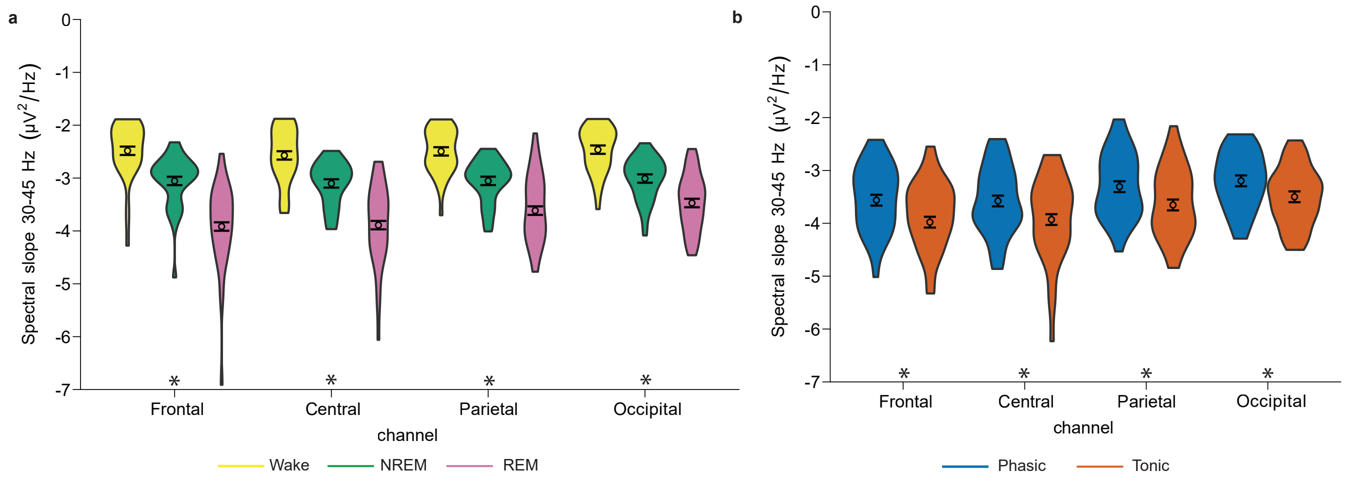


**Supplementary Figure 4**

Differences in a. wake, NREM and REM sleep and b. phasic and tonic REM sleep spectral slopes in the 30 – 45 Hz range during baseline nights averaged across frontal (F3, F4), central (C3, C4), parietal (P3, P4) and occipital (O1, O2) regions. Statistical comparisons were made using linear mixed-effects models (lme's) between wake, NREM and REM sleep (see Supplementary Table S4 for detailed models) and Tukey post-hoc tests (* $p < 0.05$ – $p < 0.001$). Violin plots show data distribution. Estimated means resulting from the lme's averaged across participants ($n = 36$) are indicated with open black circles. Error bars indicate the s.e.m.

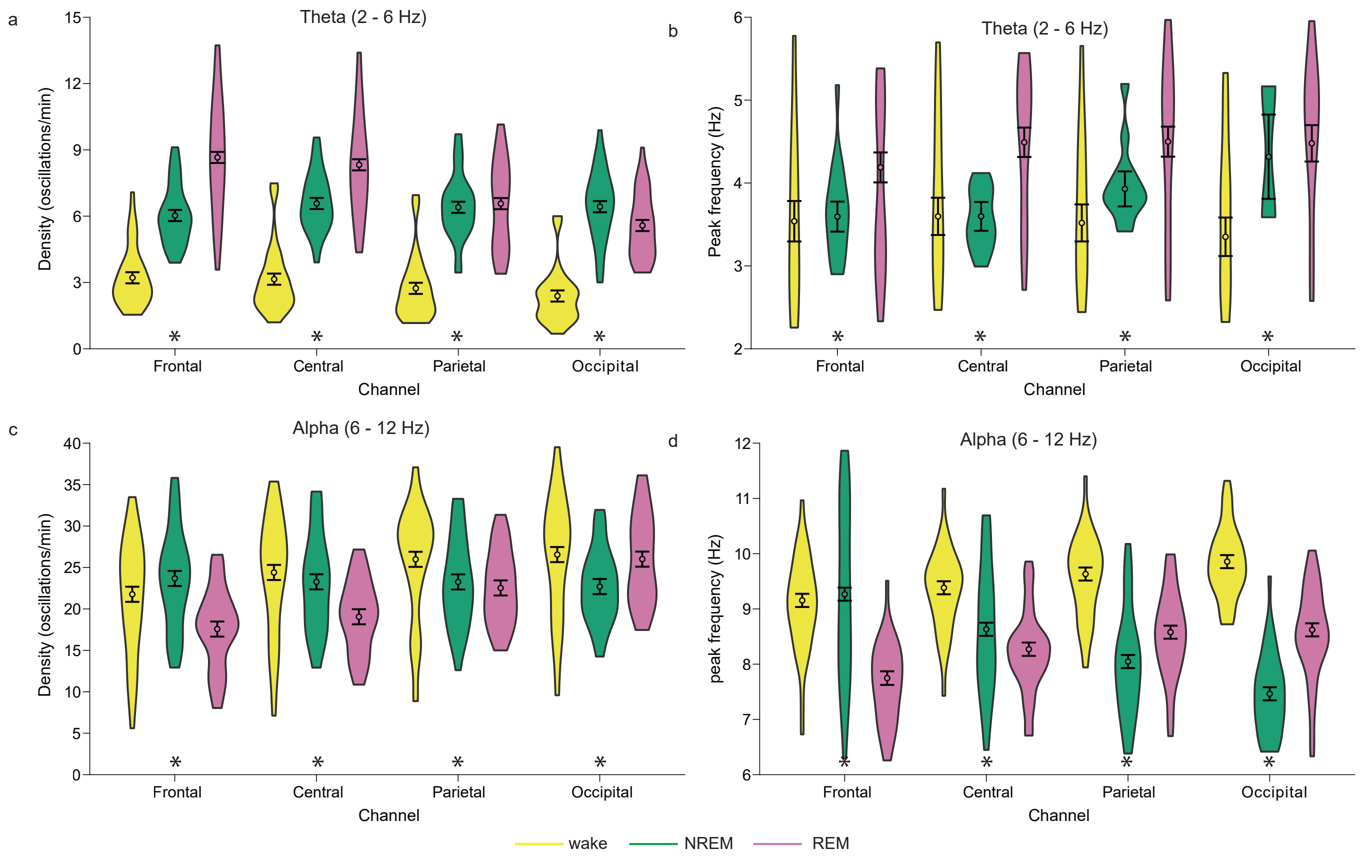


**Supplementary Figure 5**
Differences in wake, NREM and REM sleep oscillatory characteristics during baseline nights averaged across frontal (F3, F4), central (C3, C4), parietal (P3, P4) and occipital (O1, O2) regions. a-b. theta (2 - 6 Hz) oscillations' density and peak frequency. c-d. alpha (6 - 12 Hz) oscillations' density and peak frequency. Statistical comparisons were made using linear mixed-effects models (lme's) and Tukey post-hoc tests (* $p<0.05$ – $p < 0.001$) between wake, NREM and REM sleep (see Supplementary Table S4 for detailed models). Violin plots show data distribution. Estimated means resulting from the lme's averaged across participants ($n = 36$) are indicated with open black circles. Error bars indicate the s.e.m.

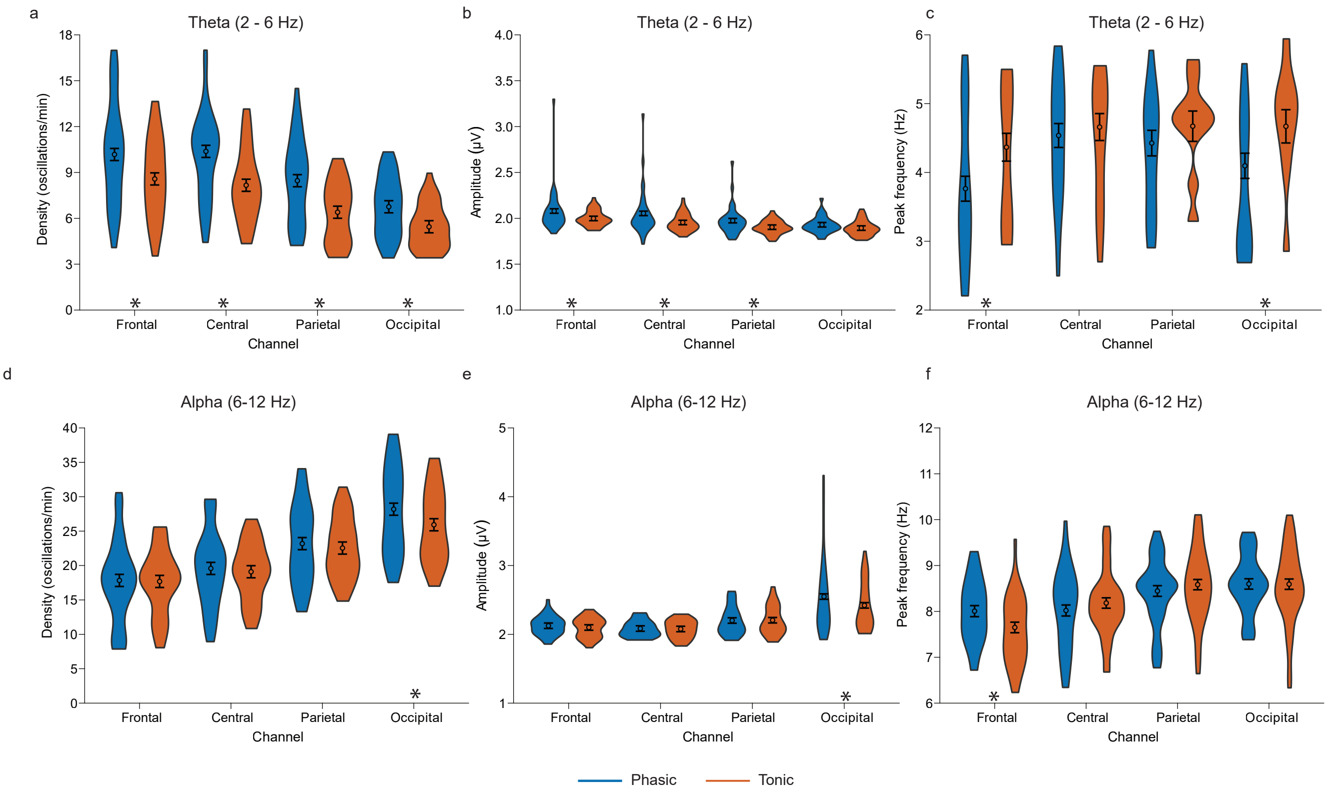


**Supplementary Figure 6**

Differences in phasic and tonic REM sleep oscillatory characteristics during baseline nights averaged across frontal (F3, F4), central (C3, C4), parietal (P3, P4) and occipital (O1, O2) regions. a-c. theta (2 - 6 Hz) oscillations' density, amplitude and peak frequency. d-f. alpha (6 - 12 Hz) oscillations' density, amplitude and peak frequency. Statistical comparisons were made using linear mixed-effects models and anova (lme's, * $p<0.05$ – $p < 0.001$) between phasic and tonic REM sleep (see Supplementary Table S4 for detailed models). Violin plots show data distribution. Estimated means resulting from the lme's averaged across participants (n = 36) are indicated with open black circles. Error bars indicate the s.e.m.

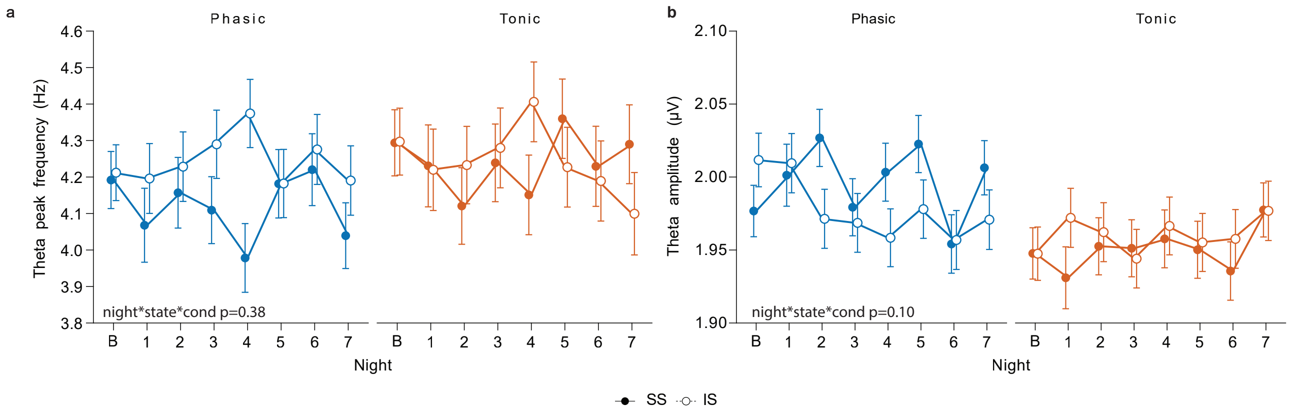


**Supplementary Figure 7**

Changes in phasic and tonic oscillatory characteristics during insufficient sleep (IS, open circles) compared to sufficient sleep (SS, filled circles). a. Changes in phasic and tonic theta oscillation peak frequency. b. Changes in phasic and tonic theta oscillation amplitude. Statistical comparisons were made using linear mixed-effects models (lme's, see Supplementary Table S3 for detailed models) and Tukey post-hoc tests (* $p<0.05$ – $p<0.001$). Data are presented as estimated means resulting from the lme's averaged across participants (n=36) for each night. Error bars indicate the standard error of the mean (s.e.m). B = baseline.

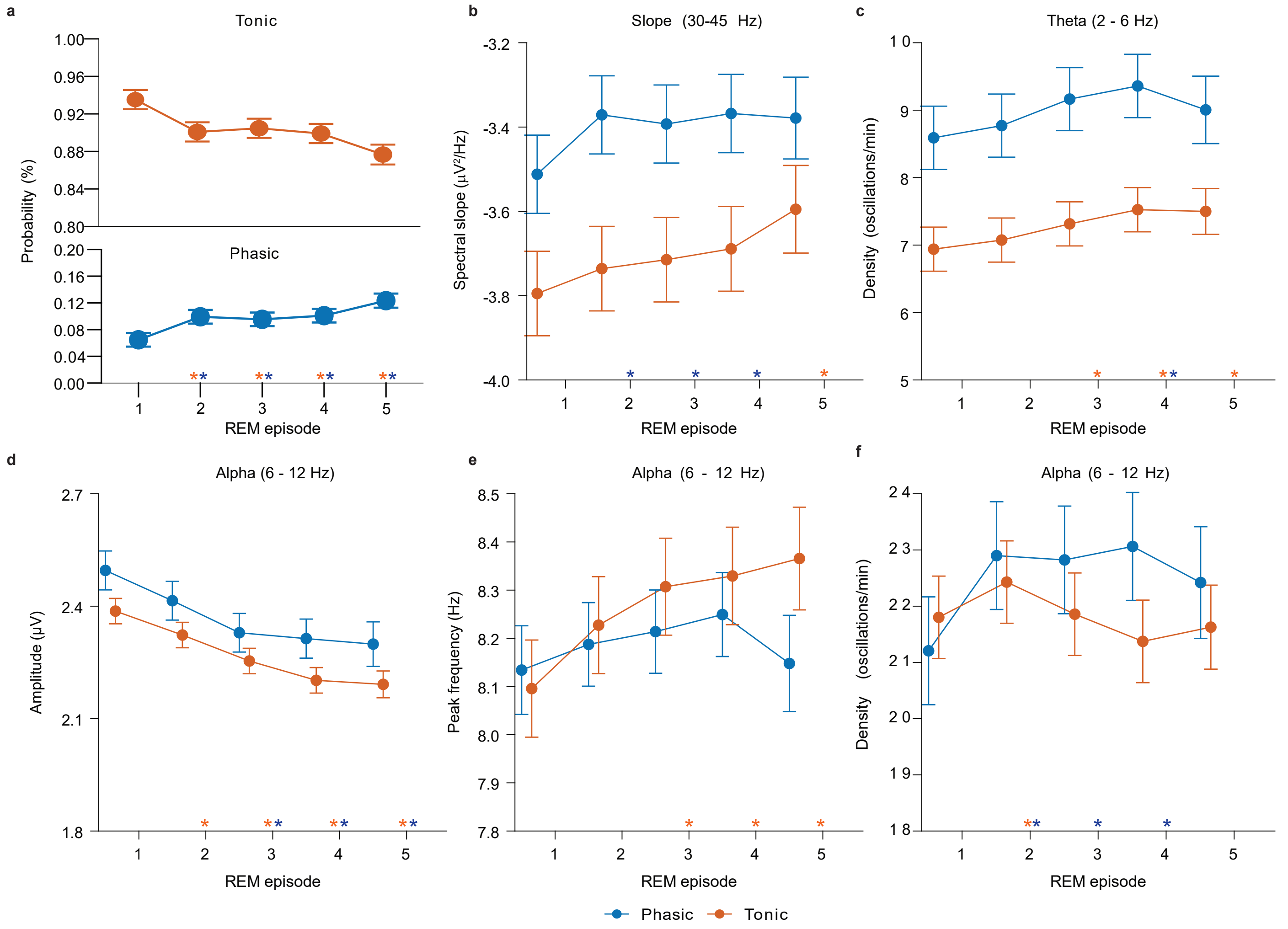


**Supplementary Figure 8**

Within-night dynamics of phasic and tonic REM oscillatory and aperiodic characteristics that changed with insufficient sleep, during baseline nights. a. probability of phasic and tonic REM sleep occurrence. b. spectral slope in the 30 – 45 Hz range c. theta oscillation density. d and e. alpha oscillation amplitude, peak frequency and density. Statistical comparisons were made using linear mixed-effects models (lme's, see Supplementary Table S4 for detailed models) and Sidak post-hoc tests in comparison to the first REM episode (* p<0.05 – p<0.001). Estimated means resulting from the lme's averaged across participants (n = 36) are indicated with coloured circles (phasic, blue; tonic, orange). Error bars indicate the s.e.m.

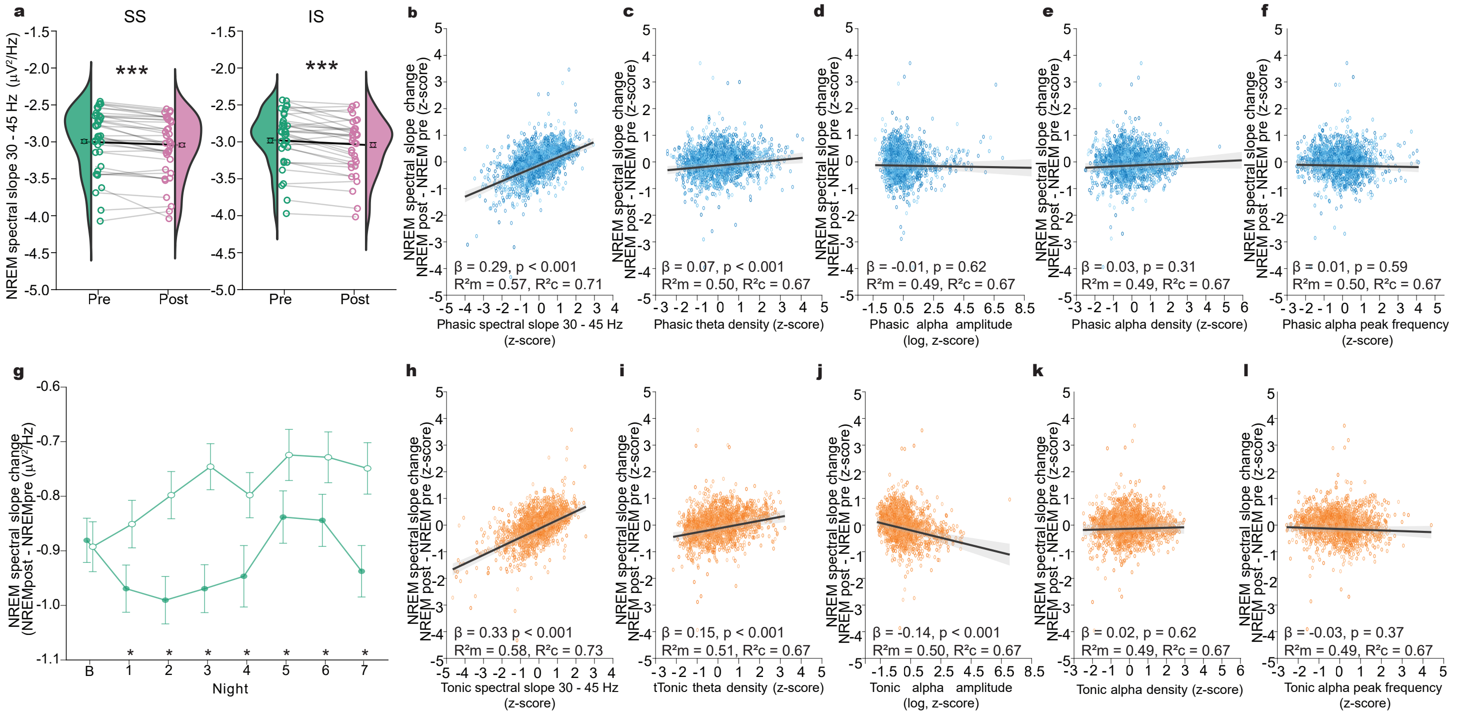


**Supplementary Figure 9**

Changes in NREM spectral slopes (30-45 Hz) from pre- to post- REM sleep period (NREM-REM-NREM triplets) and their relationship with phasic and tonic periodic and aperiodic EEG components in the REM period in-between for SS and IS nights. a. Changes in NREM spectral slopes in the 30 - 45 Hz range from before to after a REM episode (NREM-REM-NREM triplets). Statistical comparisons were made using linear mixed-effects models (lme's, data included from both SS and IS baseline nights, see Supplementary Table S4 for detailed models *** p<0.001). Coloured circles indicate average values of SS and IS baseline nights for each participant and violins indicate their distributions. Estimated means resulting from the lme's averaged across participants (n = 36) are indicated with open black circles. Error bars indicate the s.e.m. Grey and black lines connect the data points from the same participant and average, respectively. b-f. Partial linear regression plot showing relationship between NREM spectral slope change from before to after a REM episode (post-pre) with phasic spectral slope in (b), phasic theta density in (c), phasic alpha amplitude in (d), phasic alpha density in (e) and phasic alpha peak frequency in (f) for SS and IS nights. Black line represents partial linear regression with 95% confidence interval shaded in grey, orange or blue circles represent partial residuals. g. Comparison of NREM spectral slope changes from before to after a REM episode (post - pre) between SS and IS across the week. Statistical comparisons were made using linear mixed-effects models with Tukey post-hocs (lme's, see Supplementary Table S4 for detailed models * p<0.001). h-l. Same as for b-f, but for tonic REM. All correlations were FDR [False-Discovery Rate] corrected.

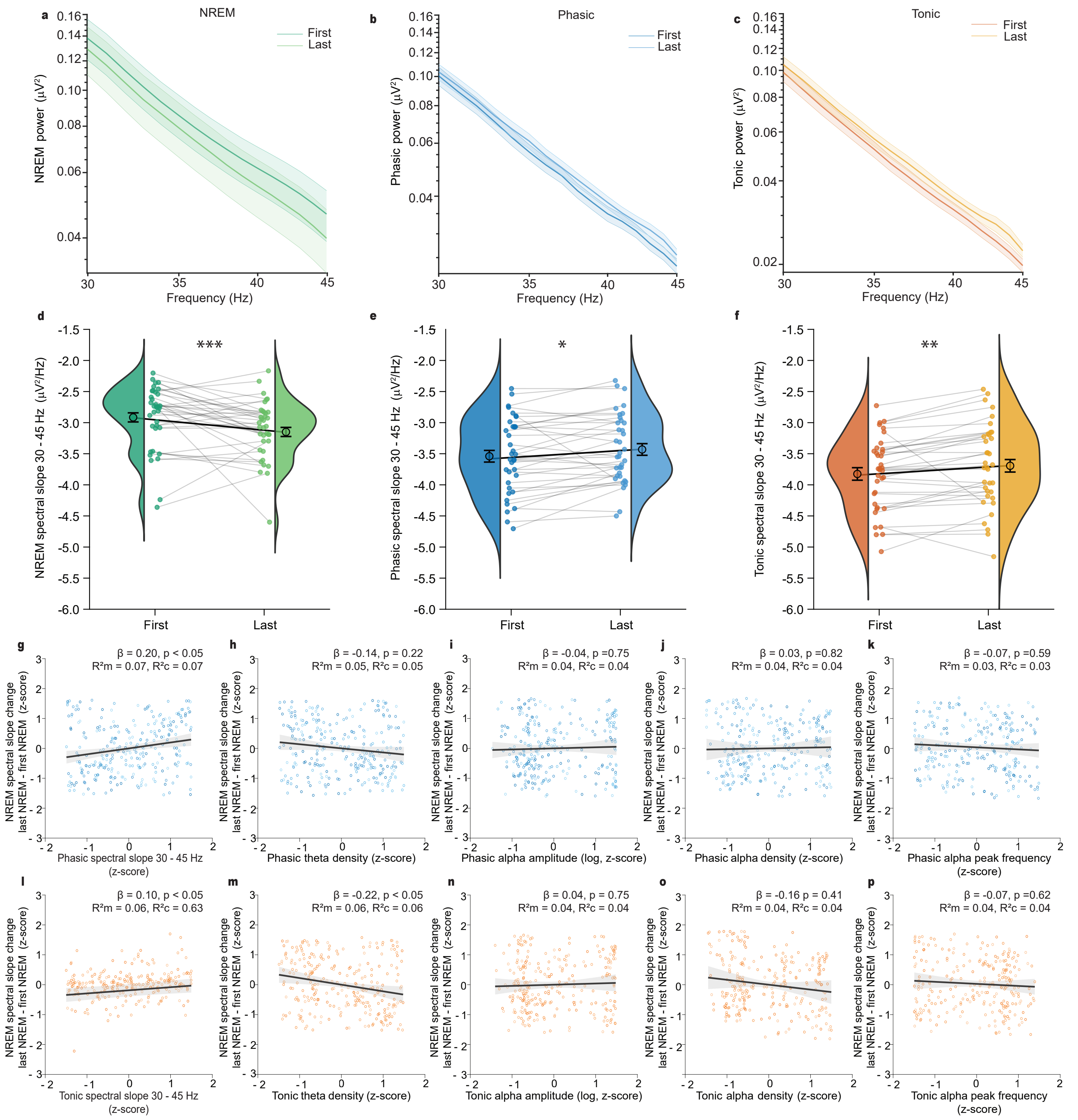


**Supplementary Figure 10**

Changes in NREM spectral slopes overnight and their relationship with phasic and tonic spectral slopes, alpha amplitude and theta density during baseline nights. a and d. Changes in NREM spectral slopes in the 30-45 Hz range from the first to the last sleep cycle, during baseline nights. b and e. Same as a and d, but for phasic REM. c and f. Same as a and d, but for tonic REM. Power spectra were averaged across participants (N = 36), electrodes, and SS and IS baseline nights. Shaded area represents the standard error of the mean (s.e.m.) across participants for the power spectra. Statistical comparisons were made using linear mixed-effects models (lme's, data included from both SS and IS baseline nights, see Supplementary Table S4 for detailed models (* $p<0.05$, ** $p<0.01$ and *** $p<0.001$). Coloured circles indicate average values of SS and IS baseline nights for each participant and violins indicate their distributions. Estimated means resulting from the lme's averaged across participants (n = 36) are indicated with open black circles. Error bars indicate the s.e.m. Grey and black lines connect the data points from the same participant and average, respectively. g. Partial linear regression plot showing relationship between NREM spectral slope change overnight with mean phasic spectral slope. h. Partial linear regression plot showing relationship between NREM spectral slope change overnight with phasic theta density. i. Partial linear regression plot showing relationship between NREM spectral slope change overnight with phasic alpha amplitude. j. Partial linear regression plot showing relationship between NREM spectral slope change overnight with phasic alpha density. k. Partial linear regression plot showing relationship between NREM spectral slope change overnight with phasic alpha peak frequency. l-p. Same as g-p, but for tonic REM. Black line represents partial linear regression with 95% confidence interval shaded in grey, orange or blue circles represent partial residuals. All correlations were FDR [False-Discovery Rate]-corrected.

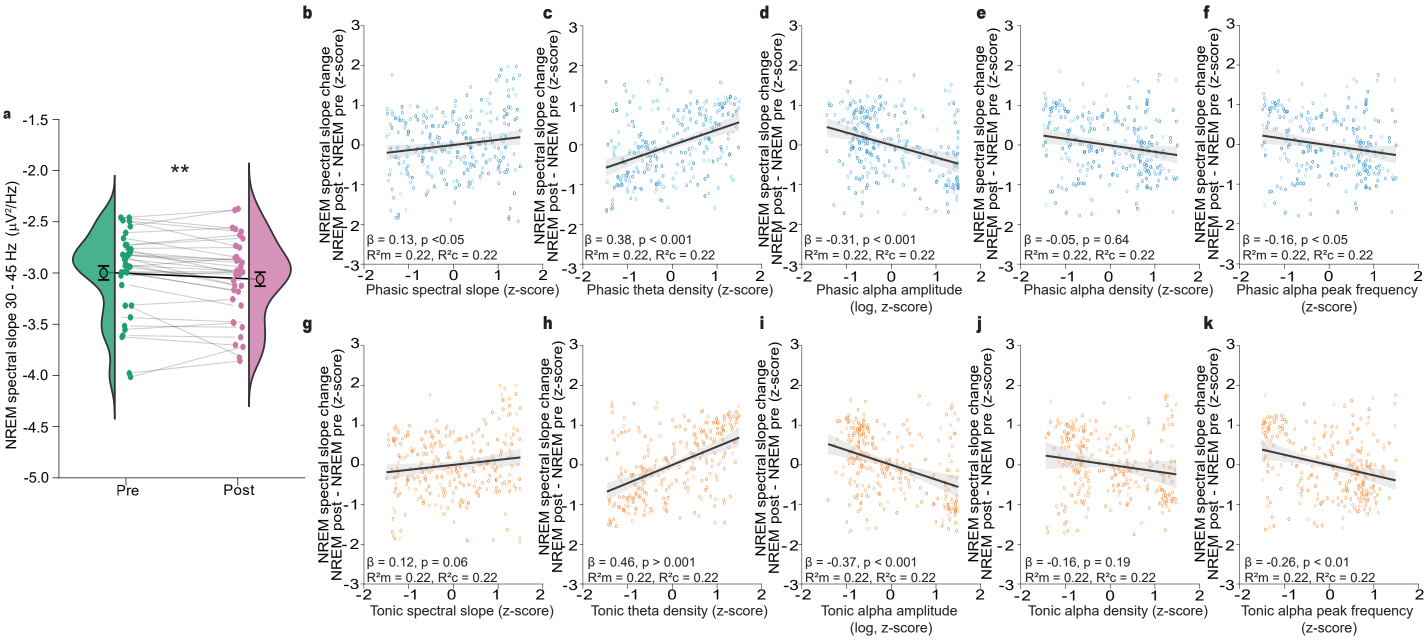


**Supplementary Figure 11**

Changes in NREM spectral slopes (30-45 Hz) from pre- to post- REM sleep period (NREM-REM-NREM triplets) and their relationship with phasic and tonic periodic and aperiodic EEG components in the REM period in-between during baseline nights. a. Changes in NREM spectral slopes in the 30 - 45 Hz range from before to after a REM episode (NREM-REM-NREM triplets).. Statistical comparisons were made using linear mixed-effects models (lme's, data included from both SS and IS baseline nights, see Supplementary Table S4 for detailed models *** p<0.001). Coloured circles indicate average values of SS and IS baseline nights for each participant and violins indicate their distributions. Estimated means resulting from the lme's averaged across participants (n = 36) are indicated with open black circles. Error bars indicate the s.e.m. Grey and black lines connect the data points from the same participant and average, respectively. b-k. Partial linear regression plot showing relationship between NREM spectral slope change from before to after a REM episode (post-pre) with phasic spectral slope in (b), phasic theta density in (c), phasic alpha amplitude in (d), phasic alpha density in (e) and phasic alpha peak frequency in (f) during baseline nights (data included from both SS and IS baseline nights). g-k. Same as for b-d but for tonic REM. Black line represents partial linear regression with 95% confidence interval shaded in grey, orange or blue circles represent partial residuals.

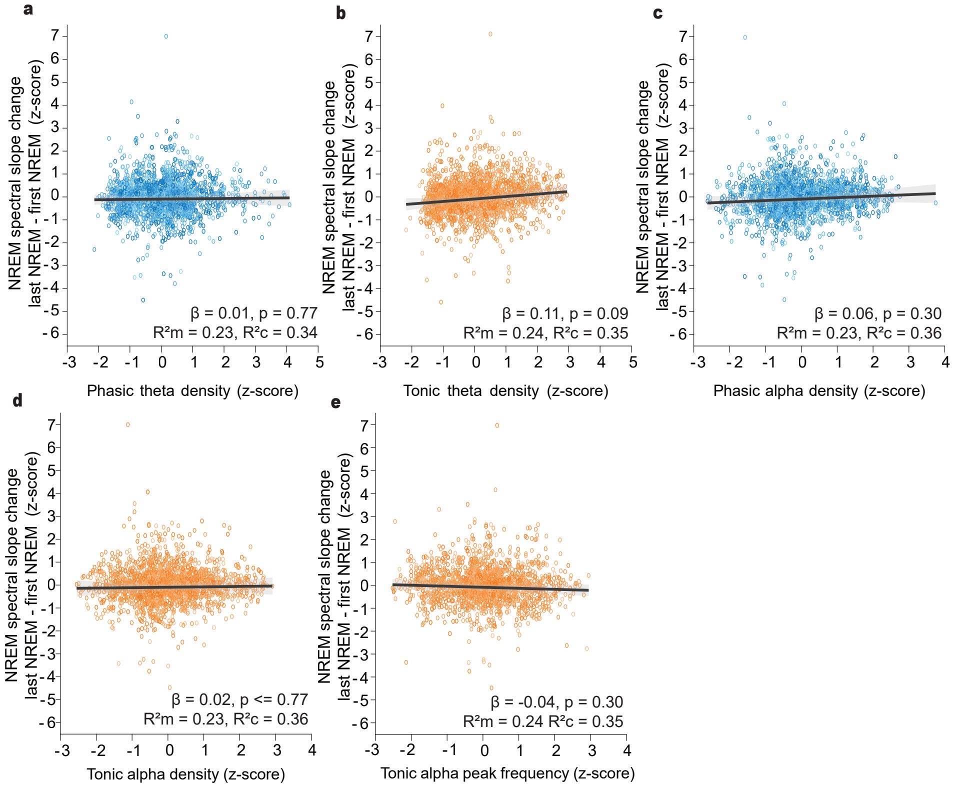


**Supplementary Figure 12**

Partial linear regression plots showing relationship between NREM spectral slope change overnight for sufficient sleep (SS) and insufficient sleep (IS) nights with (a) phasic theta density, (b) tonic theta density, (c) phasic alpha density, (d) tonic alpha density and (e) tonic alpha peak frequency. Black line represents partial linear regression with 95% confidence interval shaded in grey, orange or blue circles represent partial residuals. All correlations were FDR [False-Discovery Rate]-corrected.

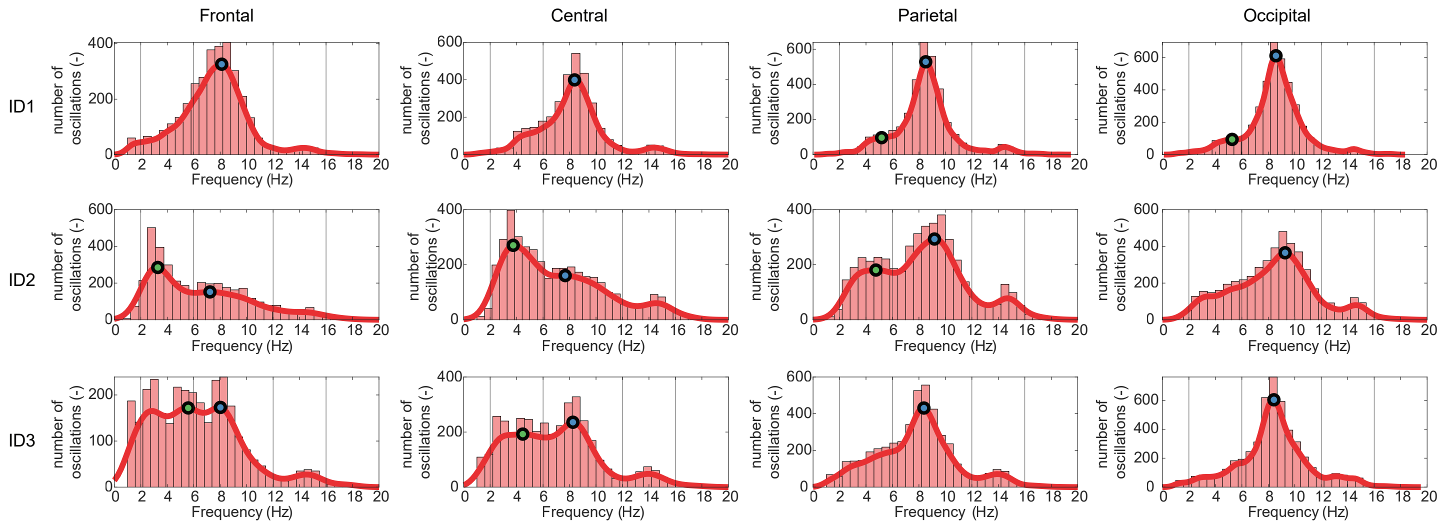


**Supplementary Figure 13**
EEG oscillations' frequency distribution histograms during REM sleep across frontal (F3), central (C3), posterior (P3) and occipital (O1) regions for 3 example participants during baseline nights.

| Night | | TST | | WASO | | N1 | | N2 | | N3 | | NREM (N2 + N3) | | phasic | | tonic | | REM | |
|---|---|---|---|---|---|---|---|---|---|---|---|---|---|---|---|---|---|---|---|
| | | Mean | S.E.M | Mean | S.E.M | Mean | S.E.M | Mean | S.E.M | Mean | S.E.M | Mean | S.E.M | Mean | S.E.M | Mean | S.E.M | Mean | S.E.M |
| **Baseline** | SS | 447.68 | 5.92 | 19.63 | 3.16 | 47.67 | 3.39 | 177.38 | 5.00 | 103.10 | 6.56 | 280.47 | 5.23 | 11.20 | 1.13 | 104.70 | 4.23 | 119.54 | 4.34 |
| | IS | 452.46 | 3.38 | 21.53 | 3.16 | 44.66 | 2.76 | 174.76 | 5.44 | 110.50 | 5.32 | 285.26 | 3.89 | 12.05 | 1.26 | 107.63 | 3.93 | 122.54 | 3.83 |
| **1** | SS | 554.66 | 5.28 | 30.93 | 4.24 | 64.03 | 3.73 | 223.97 | 6.01 | 110.66 | 5.79 | 334.63 | 4.91 | 17.16 | 1.62 | 135.63 | 4.93 | 156.00 | 4.84 |
| | IS | 341.84 | 2.84 | 12.21 | 2.52 | 31.88 | 2.01 | 116.24 | 3.79 | 102.24 | 5.53 | 218.47 | 4.24 | 7.50 | 0.81 | 81.73 | 2.91 | 91.49 | 3.05 |
| **2** | SS | 519.04 | 9.85 | 51.66 | 10.60 | 69.24 | 3.34 | 213.23 | 5.88 | 94.57 | 4.60 | 307.80 | 6.54 | 15.33 | 1.82 | 119.50 | 6.01 | 142.00 | 5.78 |
| | IS | 341.91 | 3.18 | 11.38 | 2.30 | 28.91 | 1.90 | 119.24 | 4.92 | 105.59 | 5.04 | 224.82 | 4.33 | 7.22 | 0.88 | 79.13 | 2.95 | 88.18 | 3.17 |
| **3** | SS | 515.54 | 7.76 | 65.22 | 11.20 | 66.69 | 3.55 | 216.44 | 5.08 | 94.94 | 4.95 | 311.38 | 6.64 | 16.92 | 1.58 | 116.93 | 4.51 | 137.47 | 4.69 |
| | IS | 346.63 | 1.67 | 7.93 | 1.42 | 28.16 | 1.64 | 120.01 | 5.51 | 108.04 | 5.73 | 228.06 | 3.65 | 7.28 | 0.87 | 79.95 | 3.19 | 90.41 | 3.31 |
| **4** | SS | 508.89 | 7.83 | 59.41 | 9.11 | 67.52 | 3.67 | 213.34 | 6.36 | 94.44 | 5.46 | 307.78 | 6.39 | 16.32 | 1.44 | 114.11 | 3.42 | 133.59 | 3.53 |
| | IS | 349.11 | 0.99 | 9.59 | 2.54 | 28.44 | 1.70 | 112.97 | 4.35 | 110.59 | 5.07 | 223.56 | 3.56 | 7.76 | 0.73 | 87.26 | 2.73 | 97.11 | 2.87 |
| **5** | SS | 506.52 | 9.44 | 51.70 | 8.70 | 65.92 | 3.43 | 206.77 | 6.26 | 94.02 | 5.31 | 300.79 | 7.20 | 15.62 | 1.56 | 121.18 | 4.92 | 139.80 | 5.09 |
| | IS | 345.60 | 2.59 | 7.84 | 1.43 | 29.99 | 1.87 | 115.31 | 4.23 | 105.99 | 5.36 | 221.30 | 3.65 | 7.29 | 0.82 | 85.13 | 3.03 | 94.31 | 3.12 |
| **6** | SS | 508.53 | 8.79 | 60.77 | 13.48 | 68.34 | 3.50 | 215.34 | 6.58 | 93.34 | 5.30 | 308.68 | 7.64 | 15.25 | 1.49 | 113.54 | 3.81 | 131.52 | 4.01 |
| | IS | 345.47 | 1.56 | 8.84 | 1.40 | 27.68 | 1.74 | 116.79 | 4.78 | 106.28 | 5.40 | 223.07 | 3.01 | 7.54 | 0.83 | 84.51 | 2.82 | 94.72 | 2.82 |
| **7** | SS | 451.09 | 9.90 | 94.66 | 9.79 | 57.58 | 3.42 | 183.70 | 5.48 | 95.42 | 4.37 | 279.13 | 7.26 | 12.63 | 1.23 | 97.64 | 4.23 | 114.39 | 4.49 |
| | IS | 341.16 | 2.08 | 16.26 | 2.15 | 28.65 | 1.95 | 124.65 | 5.40 | 102.66 | 5.32 | 227.31 | 3.84 | 7.97 | 0.91 | 75.72 | 3.10 | 85.21 | 3.41 |

**Supplementary Table 1**

Absolute average duration ± s.e.m. (minutes) of Total Sleep Time (TST), Wake After Sleep Onset (WASO), N1, N2, N3, NREM (N2 + N3), phasic, tonic and REM sleep during baseline and the week of sufficient (SS) and insufficient sleep (IS).

**General Positive Affect for baseline**

| | | variables | Estimate | Std. Error | df | pvalue | adj pvalue | 95% CI |
|---|---|---|---|---|---|---|---|---|
| duration | | NREM percentage | -0.0834 | 0.0085 | 9505.818 | 0 | 0 | -0.1000,-0.0667 |
| | | REM percentage | 0.0536 | 0.0082 | 9503.4105 | 0 | 0 | 0.0375, 0.0698 |
| | | phasic percentage | 0.1308 | 0.0108 | 9511.7616 | 0 | 0 | 0.1096, 0.1518 |
| | | tonic percentage | 0.037 | 0.0082 | 9503.4663 | 0 | 0 | 0.0209, 0.0532 |
| oscillatory components | phasic | IAPF | 0.0041 | 0.0102 | 7960.5009 | 0.688 | 0.881 | -0.0159, 0.0242 |
| | | alpha density | -0.0524 | 0.0144 | 9488.4597 | 3E-04 | 0.0011 | -0.0808,-0.0243 |
| | | alpha amplitude | -0.0319 | 0.0097 | 9502.1796 | 0.001 | 0.0035 | -0.0510,-0.0128 |
| | | theta density | 0.0318 | 0.0099 | 9504.3281 | 0.001 | 0.0037 | 0.0123, 0.0513 |
| | | theta amplitude | -0.0006 | 0.0069 | 9442.2388 | 0.934 | 0.9338 | -0.0142, 0.0130 |
| | tonic | IAPF | -0.0313 | 0.0113 | 9028.9948 | 0.006 | 0.0133 | -0.0535,-0.0091 |
| | | alpha density | -0.0732 | 0.016 | 9457.9565 | 0 | 0 | -0.1047,-0.0420 |
| | | alpha amplitude | -0.0529 | 0.0116 | 9509 | 0 | 0 | -0.0756,-0.0303 |
| | | theta density | 0.0704 | 0.0112 | 9508.2833 | 0 | 0 | 0.0485, 0.0923 |
| | | theta amplitude | 0.0269 | 0.0084 | 9482.2812 | 0.001 | 0.0037 | 0.0105, 0.0434 |
| spectral slope | phasic | 30 - 45 Hz | 0.002 | 0.0115 | 8692.6088 | 0.859 | 0.9338 | -0.0205, 0.0245 |
| | tonic | 30 - 45 Hz | 0.0082 | 0.012 | 8709.9776 | 0.494 | 0.6779 | -0.0154, 0.0316 |

**General Negative Affect for baseline**

| | | variables | Estimate | Std. Error | df | pvalue | adj pvalue | 95% CI |
|---|---|---|---|---|---|---|---|---|
| duration | | NREM percentage | -0.034 | 0.011 | 9500.4076 | 0.002 | 0.0047 | -0.0555,-0.0126 |
| | | REM percentage | 0.0125 | 0.0106 | 9508.1852 | 0.239 | 0.3816 | -0.0082, 0.0334 |
| | | phasic percentage | 0.0706 | 0.0139 | 9229.8035 | 0 | 0 | 0.0435, 0.0979 |
| | | tonic percentage | 0.0278 | 0.0106 | 9508.0691 | 0.009 | 0.0186 | 0.0071, 0.0486 |
| oscillatory components | phasic | IAPF | -0.0247 | 0.0135 | 7922.3917 | 0.067 | 0.1348 | -0.0511, 0.0019 |
| | | alpha density | -0.0056 | 0.0187 | 8938.7023 | 0.765 | 0.9069 | -0.0421, 0.0311 |
| | | alpha amplitude | -0.0015 | 0.0127 | 9493.5499 | 0.908 | 0.9338 | -0.0262, 0.0234 |
| | | theta density | -0.0174 | 0.0129 | 9480.5259 | 0.178 | 0.2993 | -0.0427, 0.0079 |
| | | theta amplitude | 0.0014 | 0.0091 | 9454.539 | 0.877 | 0.9338 | -0.0163, 0.0192 |
| | tonic | IAPF | -0.0097 | 0.0147 | 8882.4 | 0.508 | 0.6779 | -0.0384, 0.0192 |
| | | alpha density | 0.0302 | 0.0207 | 8553.9843 | 0.145 | 0.2574 | -0.0102, 0.0708 |
| | | alpha amplitude | -0.0258 | 0.015 | 9377.4192 | 0.085 | 0.1606 | -0.0551, 0.0037 |
| | | theta density | -0.005 | 0.0145 | 9422.9864 | 0.73 | 0.8988 | -0.0336, 0.0234 |
| | | theta amplitude | 0.0114 | 0.0109 | 9498.8144 | 0.296 | 0.4517 | -0.0100, 0.0328 |
| spectral slope | phasic | 30 - 45 Hz | -0.0015 | 0.0149 | 8444.6269 | 0.92 | 0.9338 | -0.0307, 0.0278 |
| | tonic | 30 - 45 Hz | -0.0114 | 0.0156 | 8383.0717 | 0.465 | 0.6758 | -0.0420, 0.0192 |

**Supplementary Table 2**

Relationship of general positive and negative affect with NREM, REM, phasic and tonic proportion, oscillatory and aperiodic characteristics during baseline nights assessed using lme's (n=36; data included from both SS and IS, see Supplementary Table S4 for detailed models).

**Dimension 1 for baseline**

| | | variables | Estimate | Std. Error | df | pvalue | adj pvalue | 95% CI |
|---|---|---|---|---|---|---|---|---|
| duration | | NREM percentage | -0.0998 | 0.0307 | 1405.5084 | 0.0012 | 0.0059 | -0.1598,-0.0397 |
| | | REM percentage | -0.0438 | 0.028 | 1412.1817 | 0.1187 | 0.2532 | -0.0983, 0.0117 |
| | | phasic percentage | -0.4485 | 0.0429 | 1353.4151 | 0 | 0 | -0.5322,-0.3604 |
| | | tonic percentage | 0.0066 | 0.0293 | 1408.9397 | 0.8223 | 0.8771 | -0.0504, 0.0642 |
| oscillatory components | phasic | IAPF | -0.0245 | 0.0274 | 1155.8637 | 0.3721 | 0.5954 | -0.0777, 0.0299 |
| | | alpha density | -0.0538 | 0.0387 | 1389.9663 | 0.1649 | 0.3198 | -0.1290, 0.0224 |
| | | alpha amplitude | 0.0073 | 0.0257 | 1401.6242 | 0.7768 | 0.8572 | -0.0429, 0.0576 |
| | | theta density | 0.0105 | 0.0264 | 1405.5944 | 0.6917 | 0.8115 | -0.0413, 0.0621 |
| | | theta amplitude | 0.0775 | 0.0215 | 1385.7095 | 0.0003 | 0.0016 | 0.0355, 0.1195 |
| | tonic | IAPF | -0.0348 | 0.0301 | 1348.9957 | 0.2478 | 0.4531 | -0.0932, 0.0245 |
| | | alpha density | 0.034 | 0.0436 | 1353.1668 | 0.436 | 0.6489 | -0.0508, 0.1196 |
| | | alpha amplitude | -0.0331 | 0.0298 | 1408.4016 | 0.2669 | 0.4617 | -0.0910, 0.0255 |
| | | theta density | 0.0134 | 0.0284 | 1408.1725 | 0.6369 | 0.7839 | -0.0422, 0.0688 |
| | | theta amplitude | 0.0037 | 0.0224 | 1393.6421 | 0.8692 | 0.883 | -0.0399, 0.0476 |
| spectral slope | phasic | 30 - 45 Hz | -0.0372 | 0.0278 | 1318.9478 | 0.1808 | 0.3403 | -0.0914, 0.0173 |
| | tonic | 30 - 45 Hz | 0.0212 | 0.0284 | 1323.9253 | 0.4548 | 0.651 | -0.0344, 0.0766 |

**Dimension 2 for baseline**

| | | variables | Estimate | Std. Error | df | pvalue | adj pvalue | 95% CI |
|---|---|---|---|---|---|---|---|---|
| duration | | NREM percentage | 0.0267 | 0.0247 | 1411.4925 | 0.2801 | 0.4717 | -0.0215, 0.0751 |
| | | REM percentage | -0.0541 | 0.0224 | 1408.4257 | 0.0158 | 0.0632 | -0.0981,-0.0104 |
| | | phasic percentage | 0.2234 | 0.0352 | 1385.8422 | 0 | 0 | 0.1539, 0.2919 |
| | | tonic percentage | -0.1021 | 0.0233 | 1410.1635 | 0 | 0 | -0.1476,-0.0566 |
| oscillatory components | phasic | IAPF | 0.0604 | 0.0214 | 1144.5368 | 0.0048 | 0.0219 | 0.0186, 0.1022 |
| | | alpha density | 0.1202 | 0.0309 | 1408.5214 | 0.0001 | 0.0006 | 0.0596, 0.1803 |
| | | alpha amplitude | -0.0038 | 0.0205 | 1390.7307 | 0.8541 | 0.8817 | -0.0437, 0.0363 |
| | | theta density | 0.0551 | 0.021 | 1394.6233 | 0.0089 | 0.038 | 0.0141, 0.0963 |
| | | theta amplitude | -0.1678 | 0.0164 | 1375.3839 | 0 | 0 | -0.1997,-0.1358 |
| | tonic | IAPF | 0.0085 | 0.0239 | 1341.2329 | 0.7234 | 0.8122 | -0.0380, 0.0555 |
| | | alpha density | -0.0139 | 0.0351 | 1407.8554 | 0.693 | 0.8115 | -0.0822, 0.0549 |
| | | alpha amplitude | 0.0241 | 0.0237 | 1398.7088 | 0.311 | 0.5104 | -0.0221, 0.0709 |
| | | theta density | -0.0129 | 0.0226 | 1397.9051 | 0.5693 | 0.7752 | -0.0570, 0.0315 |
| | | theta amplitude | -0.0303 | 0.0177 | 1385.0997 | 0.088 | 0.2011 | -0.0649, 0.0045 |
| spectral slope | phasic | 30 - 45 Hz | 0.0487 | 0.0224 | 1310.3154 | 0.0296 | 0.0902 | 0.0047, 0.0922 |
| | tonic | 30 - 45 Hz | -0.009 | 0.0232 | 1317.434 | 0.6974 | 0.8115 | -0.0546, 0.0361 |

**Dimension 3 for baseline**

| | | variables | Estimate | Std. Error | df | pvalue | adj pvalue | 95% CI |
|---|---|---|---|---|---|---|---|---|
| duration | | NREM percentage | -0.0494 | 0.026 | 1412.3841 | 0.0574 | 0.1531 | -0.0999, 0.0024 |
| | | REM percentage | -0.0176 | 0.0236 | 1410.6146 | 0.4577 | 0.651 | -0.0643, 0.0285 |
| | | phasic percentage | -0.1496 | 0.0373 | 1370.6679 | 0.0001 | 0.0006 | -0.2221,-0.0750 |
| | | tonic percentage | -0.0415 | 0.0247 | 1412.2189 | 0.0924 | 0.2039 | -0.0903, 0.0064 |
| oscillatory components | phasic | IAPF | -0.0418 | 0.0226 | 1147.9324 | 0.0642 | 0.1644 | -0.0859, 0.0023 |
| | | alpha density | -0.0779 | 0.0327 | 1408.9777 | 0.0173 | 0.0651 | -0.1413,-0.0127 |
| | | alpha amplitude | 0.0171 | 0.0216 | 1393.5791 | 0.4279 | 0.6489 | -0.0249, 0.0597 |
| | | theta density | 0.0316 | 0.0222 | 1397.1927 | 0.1549 | 0.3098 | -0.0120, 0.0749 |
| | | theta amplitude | 0.1413 | 0.0176 | 1377.2189 | 0 | 0 | 0.1067, 0.1756 |
| | tonic | IAPF | -0.0125 | 0.0251 | 1343.8908 | 0.6196 | 0.7839 | -0.0614, 0.0370 |
| | | alpha density | -0.0177 | 0.037 | 1403.5587 | 0.6317 | 0.7839 | -0.0895, 0.0562 |
| | | alpha amplitude | 0.0092 | 0.0251 | 1401.7268 | 0.7138 | 0.8122 | -0.0396, 0.0586 |
| | | theta density | 0.0028 | 0.0239 | 1401.1028 | 0.9067 | 0.9067 | -0.0441, 0.0494 |
| | | theta amplitude | 0.0037 | 0.0188 | 1387.226 | 0.8455 | 0.8817 | -0.0331, 0.0403 |
| spectral slope | phasic | 30 - 45 Hz | -0.013 | 0.0237 | 1313.9555 | 0.5843 | 0.7791 | -0.0592, 0.0333 |
| | tonic | 30 - 45 Hz | 0.0519 | 0.0243 | 1320.5166 | 0.0325 | 0.0945 | 0.0046, 0.0994 |

**Dimension 4 for baseline**

| | | variables | Estimate | Std. Error | df | pvalue | adj pvalue | 95% CI |
|---|---|---|---|---|---|---|---|---|
| duration | | NREM percentage | -0.1324 | 0.0358 | 1350.7406 | 0.0002 | 0.0012 | -0.2019,-0.0619 |
| | | REM percentage | 0.3099 | 0.0318 | 1399.292 | 0 | 0 | 0.2466, 0.3717 |
| | | phasic percentage | 0.6163 | 0.0486 | 1118.6907 | 0 | 0 | 0.5158, 0.7115 |
| | | tonic percentage | 0.2761 | 0.0334 | 1390.5698 | 0 | 0 | 0.2093, 0.3411 |
| oscillatory components | phasic | IAPF | -0.0082 | 0.0332 | 1154.7057 | 0.8041 | 0.8722 | -0.0735, 0.0563 |
| | | alpha density | 0.0875 | 0.0444 | 1302.4303 | 0.0491 | 0.1366 | 0.0013, 0.1752 |
| | | alpha amplitude | 0.015 | 0.0298 | 1408.023 | 0.614 | 0.7839 | -0.0432, 0.0734 |
| | | theta density | -0.0717 | 0.0306 | 1408.607 | 0.0191 | 0.0667 | -0.1321,-0.0123 |
| | | theta amplitude | -0.055 | 0.025 | 1392.9847 | 0.0278 | 0.089 | -0.1040,-0.0064 |
| | tonic | IAPF | 0.0278 | 0.0348 | 1337.5758 | 0.4244 | 0.6489 | -0.0404, 0.0956 |
| | | alpha density | 0.0567 | 0.05 | 1214.0775 | 0.2567 | 0.4564 | -0.0402, 0.1552 |
| | | alpha amplitude | 0.0615 | 0.0344 | 1404.8284 | 0.0743 | 0.1829 | -0.0060, 0.1286 |
| | | theta density | -0.0766 | 0.0328 | 1404.7193 | 0.0198 | 0.0667 | -0.1414,-0.0129 |
| | | theta amplitude | -0.0457 | 0.0259 | 1400.7619 | 0.0784 | 0.1858 | -0.0966, 0.0049 |
| spectral slope | phasic | 30 - 45 Hz | 0.0477 | 0.0327 | 1302.4748 | 0.1452 | 0.2998 | -0.0157, 0.1126 |
| | tonic | 30 - 45 Hz | 0.0213 | 0.0334 | 1302.1455 | 0.5235 | 0.7283 | -0.0434, 0.0880 |

**Supplementary Table 3**
Relationship of principal components (PCs) of cognition with phasic and tonic duration, oscillatory and aperiodic characteristics during baseline nights assessed using lme's (n=36; data included from both SS and IS, see Supplementary Table S4 for detailed models).

**List of linear mixed-effects models:**
**Duration: Fig 1 and S2**
For the effects of SS/IS on change in duration for each vigilance state, the following model was used:
duration ~ night + night:condition + baseline duration value + order (first extension or restriction) + 1|participant

**Oscillatory and slope components: Figure 2, S3-7**
For oscillatory and slope characteristics (known here as "variable") differences between vigilant states prior sleep interventions, the following model was used with only the baseline nights:
variable ~ state*region + total sleep time + order + 1|participant
For the effects of SS/IS on oscillatory and slope characteristics, the following model was used:
variable ~night*state + night:condition + state:night:condition + condition:region + region + baseline variable value + order + total sleep time + (1|participant)

**PANAS and PCA: Figure 4 and 5**

For the effects of SS/IS on change in PANAS and PCA components, the following model was used. PANAS and PCA components were z-scored normalised and rank-transformed (PANAS only) to achieve normal distribution of residuals.
PANAS/PCA score ~ condition:day + baseline variable score + day + time of test + order + (1|participant)

For associations between oscillatory and slope characteristics (variable) with PANAS and PCA component changes, models were conducted separately for phasic and tonic REM:
**Baseline:**
for sleep stage durations: PANAS/PCA score ~ variable + time of test + order + (1|participant)
for periodic/aperiodic characteristics: PANAS/PCA score ~ variable + time of test + order + channel + total sleep time + (1|participant)
**SS/IS week:**
for sleep stage durations: PANAS/PCA score ~ variable + variable baseline value + PANAS/PCA baseline value + condition:day + day + time of test + order + (1|participant)
for periodic/aperiodic characteristics: PANAS/PCA score ~ variable + variable baseline value + PANAS/PCA baseline value + condition:day + day + time of test + order + total sleep time + channel + (1|participant)

**Excitability changes overnight and across NREM-REM-NREM triplets: Figure 6, S9-12**
For changes in overnight NREM slope activity and across triplets during baseline or SS/IS:
**Baseline (S10d-f):**
NREM/phasic/tonic slope value ~ time(first/last or pre/post spectral slope value) + order + region + total sleep time + 1|participant
**SS/IS week (Fig6) :**
Figure 6a,b,d,e,g,h: NREM/phasic/tonic slope value ~ night*time + region + baseline slope value + total sleep time + order + (1|participant). Model was conducted separated for SS or IS.
Figure 6c,f,i: NREM/phasic/tonic slope change ~ night + region + night:condition + condition:region + baseline NREM/phasic/tonic slope change value + total sleep time + order + (1|participant)

For associations between slope and oscillation components with overnight NREM slope activity, models were conducted separately for phasic and tonic REM :
**Baseline:**
NREM slope change ~ slope/oscillation in-between average value + region + total sleep time + order + (1|participant)
**SS/IS week:**
NREM slope change ~ slope/oscillation in-between average value + NREM baseline slope change value + baseline slope/oscillation slope value + night:condition + night + region + total sleep time + order + (1|participant)

**Baseline night within-night dynamics: Figure S8**
To investigate variable dynamics within the night:
variable ~ cycle*region + condition + total sleep time + (1|participant)

### Supplementary Table 4
List of all linear mixed-effect models (lme's) used for statistical assessment.
